# A drug repurposing screen identifies altiratinib as a selective inhibitor of a key regulatory splicing kinase and a potential therapeutic for toxoplasmosis and malaria

**DOI:** 10.1101/2021.11.03.467097

**Authors:** Christopher Swale, Valeria Bellini, Matthew W. Bowler, Nardella Flore, Marie-Pierre Brenier-Pinchart, Dominique Cannella, Lucid Belmudes, Caroline Mas, Yohann Couté, Fabrice Laurent, Artur Scherf, Alexandre Bougdour, Mohamed-Ali Hakimi

## Abstract

The apicomplexa comprise a large phylum of single-celled, obligate intracellular protozoa that infect humans and animals and cause severe parasitic diseases. Available therapeutics against these devastating diseases are limited by suboptimal efficacy and frequent side effects, as well as the emergence and spread of resistance. Here, we use a drug repurposing strategy and identify altiratinib, a compound originally developed to treat glioblastoma, as a promising drug candidate with broad spectrum activity against apicomplexans. Altiratinib is parasiticidal and blocks the development of intracellular zoites in the nanomolar range and with a high selectivity index. We have identified *Tg*PRP4K of *T. gondii* as the primary target of altiratinib by genetic target deconvolution, highlighting key residues within the kinase catalytic site that, when mutated, confer resistance to the drug. We have further elucidated the molecular basis of the inhibitory mechanism and species selectivity of altiratinib for *Tg*PRP4K as well as for its *P. falciparum* counterpart *Pf*CLK3. Our data also point to structural features critical for binding of the other *Pf*CLK3 inhibitor, TCMDC-135051. Consistent with the role of this kinase family in splicing in a broad spectrum of eukaryotes, we have shown that altiratinib causes global disruption of splicing, primarily through intron retention in both *T. gondii* and *P. falciparum*. Thus, our data establish parasitic PRP4K/CLK3 as a promising pan-apicomplexan target whose repertoire of inhibitors can be expanded by the addition of altiratinib.

## Introduction

Infectious diseases caused by apicomplexan parasites remain the leading cause of morbidity and mortality around the world, but with even more devastating consequences in low-income countries, underscoring the need for effective medicines (De Rycker *et al.,* 2018). Indeed, *Plasmodium falciparum* causes malaria in over 200 million people worldwide and is responsible for more than 405 000 deaths in 2019 (WHO, World Malaria Report). Similarly, *Toxoplasma gondii*, the causative agent of toxoplasmosis, causes widespread zoonotic infection, with nearly one-third of the world’s population being seropositive for this parasite. In healthy adults, the acute infection resolves rapidly, leaving a chronic, subclinical infection. However, in the absence of sustained immunity, reactivation of latent forms of *T. gondii* leads to severe, life-threatening disease, as has been observed in AIDS, organ transplant or chemotherapy patients, with a high mortality rate if no treatment is given (Montoya and Liesenfeld, 2004). More severe cases may also occur following congenital transmission of the parasite to the unborn child. In addition, *Toxoplasma gondii*, together with other coccidian parasites, e.g. *Eimeria* spp. and *Neospora caninum*, are of veterinary importance as they cause significant economic losses in livestock.

For many of these apicomplexa-mediated diseases, current treatments are suboptimal, and for some there are few, if any, alternatives. Indeed, current standard treatment for toxoplasmosis is hampered by severe side effects, particularly in immunocompromised individuals (Dunay *et al.,* 2018). For malaria, artemisinin-based combination therapies (ACT) are currently used as first-line treatments in endemic countries worldwide, but the emergence and spread of resistance not only to artemisinin but also to the drug combinations is a growing threat (De Rycker *et al.,* 2018). The frequent side effects and the ever-present threat of drug resistance have led to the search for other therapeutic alternatives. Older drugs have recently made a comeback by being repurposed for new diseases to accelerate drug development. After phenotypic screening for drug repurposing, new indications for existing drugs can be quickly identified and clinical trials can be rapidly conducted. Identifying the target and understanding the mechanism of action is a critical bottleneck in drug development. Recent advances in genomics and target deconvolution strategies have shifted the problem to a plethora of putative targets awaiting clarification.

Here, we report the identification of altiratinib from a library of approved drugs that exhibits potent, nanomolar, broad-spectrum anti-apicomplexan activity with a high selectivity index. Altiratinib was in phase 1 clinical development for the treatment of invasive solid tumors, including glioblastoma (Kwon Y et al. 2015; Smith DB et al. 2015). Using a genetic target-deconvolution strategy, we identified *T. gondii Tg*PRP4K, the closest relative of the human splicing factor kinase PRP4 kinase (PRP4K/PRPF4B) and *Plasmodium falciparum Pf*CLK3 (Alam MM et al. 2019; Mahindra A et al. 2020), as the primary target of altiratinib. Using an integrated structural biology approach, we further elucidated the molecular basis for the mechanism of inhibition of altiratinib and the remarkable selectivity for the parasitic PRP4K*/*CLK3 enzymes. This kinase family plays a critical role in cell cycle progression by regulating pre-mRNA splicing in all eukaryotic lineages (Schneider M et al. 2010; Lützelberger M and Käufer NF, 2012; Corkery DP et al. 2015; Eckert D et al. 2016). Accordingly, altiratinib causes global disruption of splicing with exon skipping, intron retention, and premature transcription termination in both *T. gondii* and *P. falciparum,* but not in *Cryptosporidium parvum*, in which the kinase has significantly divergent variations that may result in resistance to altiratinib. Overall, our findings support this family of parasitic kinases as a promising apicomplexan target and highlight the structural determinants that explain the remarkable selectivity of altiratinib.

## Results

### A drug repurposing screen identifies altiratinib as a potent and selective apicomplexan inhibitor of parasite growth

To identify new drug candidates against toxoplasmosis and potential targets, we screened a small library of approved drugs for their ability to inhibit tachyzoite growth. All compounds are structurally diverse, cell permeable, medically active, and commercially available (Supplementary Table 1). Screening was performed in duplicate at 5 μM while pyrimethamine was used as a reference drug and blocked the growth of parasites as expected. The compounds that showed reproducible inhibition of parasite growth of >70% were selected for further testing (Extended Data Fig. 1a). Of the 432 compounds in the collection, 84 primary hits were found to inhibit parasite growth without detectable cytotoxicity to the host cell (Fig. 1a), preferentially targeting the cell cycle and tyrosine kinase/adaptor signaling pathways (Extended Data Fig. 1b; Supplementary Table 1). A second screen at 1 μM of the 84 compounds identified 14 molecules with EC_50_ in the nM range (Fig. 1b). The most potent growth inhibitor we identified was altiratinib (DCC-2701, DP -5164, Fig. 1c) with an EC_50_ of 28 nM against tachyzoites, which is 11-fold lower than pyrimethamine (300 nM), the standard treatment for toxoplasmosis (Fig. 1d). Altiratinib-treated parasites were smaller than the control group and no longer divided, as no daughter cells were detectable (Fig. 1e). Plaque assays showed sustained inhibition of parasite growth, as plaques could no longer be detected in the presence of altiratinib, suggesting a defect in one or more steps of the lytic cycle (Extended Data Fig. 1d). Interestingly, we did not observe regrowth after discontinuation of altiratinib, suggesting that the drug has a cidal effect in contrast to pyrimethamine (Extended Data Fig. 1d). Remarkably, altiratinib showed low host cytotoxicity, resulting in a high selectivity index (SI) with a value of 400 for human primary fibroblasts (Fig. 1f; Extended Data Fig. 1c). Altiratinib is also effective in inhibiting the growth of coccidial parasites of veterinary importance such as *Eimeria tenella* (Fig. 1g) and *Neospora caninum* (Fig. 1h), as well as *P. falciparum*, although its efficacy is lower compared with the antimalarial drug dihydroartemisinin (DHA) (Fig. 1i).

**Fig. 1.**
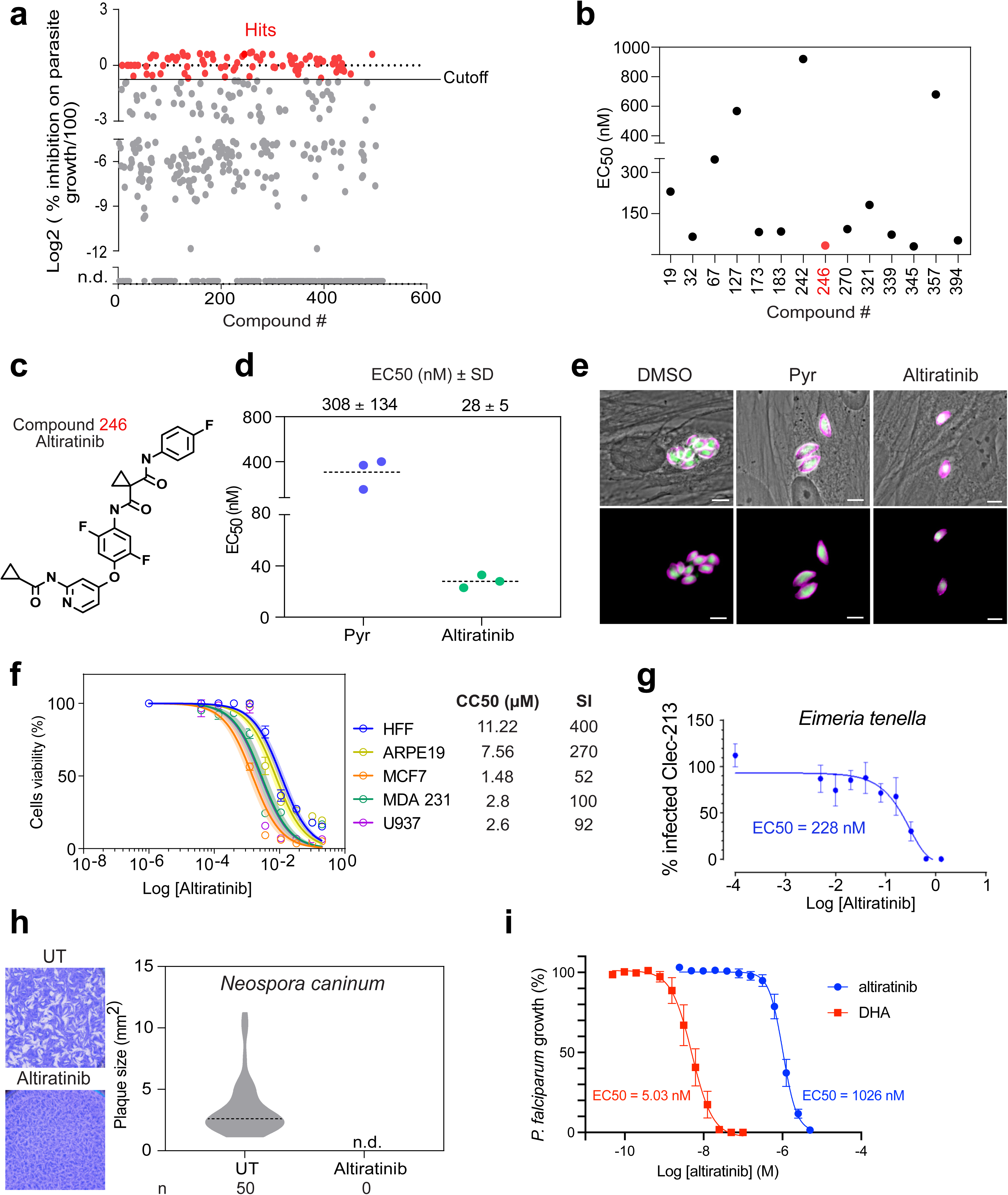
Efficacy of altiratinib against the parasite *Toxoplasma gondii*. **a,** Graphical representation of data from the medium-throughput screen. A cutoff was set at 70% of parasite inhibition. Red dots, hits. The workflow used for the screening is shown in **Extended Data** Fig. 1a. **b,** The half-maximal effective concentration (EC_50_) values of the 14 molecules validated at 1µM. **c,** Chemical structure of altiratinib. **d,** EC_50_ values for pyrimethamine and altiratinib. The confluent HFF monolayer was infected with tachyzoites of the *T. gondii* RH_NanoLucEmGFP strain (Supplementary Table 5). The EC_50_ values of each biological replicate were determined by non-linear regression analysis. EC_50_ data are presented as mean G SD from 3 independent biological replicates, each with 3 technical replicates. **e,** Compound efficiency presented by IFA. Confluent HFFs were infected with *T. gondii* RH_NanoLucEmGFP and incubated with 1µM of pyrimethamine, 300 nM of altiratinib or 0.1% of the vehicle (DMSO) for 24h. Fixed cells were stained with anti-inner membrane complex protein (GAP45) antibody (magenta). In green the cytosolic GFP. Scale bar corresponds to 5µm. **f,** Dose-response curves of HFFs, ARPE-19, MCF7, MDA231 and U937 cell lines in the presence of altiratinib. Human cells were plated out and incubated with increasing concentrations of the drug. After 72h, cell viability was determined using the “*CellTiter-Blue Assay”* kit (Promega) and cell cytotoxicity concentration (CC50) was calculated. The graph is representative of two different experiments performed in triplicate. The shaded error envelopes indicate 95% confidence intervals. On the right, CC50 values show the mean of two experiments. Selectivity index (SI) is based on the average of human CC50 divided by the average of *T. gondii* EC50. **g,** Effect concentration curve of *Eimeria tenella* in presence of altiratinib. **h,** Altiratinib inhibition of *Neospora caninum* proliferation shown by plaque assay. After 7 days of infection and drugs incubation, the size of at least 50 plaques were measured. n.d., not detected. **i,** Dose-response curves of altiratinib and dihydroartemisinin (DHA) in *P. falciparum* asexual blood-stage. Graph is representing the mean and SD values obtained in three independent experiments run in triplicate.

### Altiratinib target deconvolution by EMS-based mutagenesis-based forward genetic screen

Altiratinib was originally identified to inhibit tumor growth and invasion in a bevacizumab-resistant glioblastoma mouse model and was in phase 1 clinical development for the treatment of invasive solid tumors. The drug was predicted to be a pan-tyrosine kinase inhibitor of MET, TIE2, VEGFR2, and TRK (Kwon Y et al. 2015; Smith DB et al. 2015), but none of these kinases are conserved in apicomplexa. Therefore, to explore the mechanism of action of altiratinib in *T. gondii*, we performed a forward genetic screen combining chemical mutagenesis and RNA sequencing, as previously described (Bellini *et al.,* 2020) (Extended Data Fig. 1e). Altiratinib-resistant parasites were generated in 6 independent chemical mutagenesis experiments using 7 mM ethyl methanesulfonate (EMS) followed by selection in the presence of 300 nM altiratinib, i.e. 10-fold the EC_50_ value, for approximately 4 weeks. The resistant parasite lines were then cloned by limited dilution and a single clone from each mutagenesis experiment (designated A to F) was analyzed by whole-genome RNA sequencing (RNA-Seq). To map the EMS-induced mutations conferring resistance to altiratinib, Illumina sequencing reads were aligned to the *T. gondii GT1* reference genome. Using the parental strain as a reference, single nucleotide variations (SNVs) were identified in the assembled sequences of the resistant mutants (see *Materials and Methods*). By focusing on mutations in coding sequences, a single gene, *TGGT1_313180*, contained SNVs that resulted in amino acid changes (F647S, L686F, L715F) not present in the parental strain in five of the six drug-resistant lines (Fig. 2a,b and Supplementary Table 2).

**Fig. 2.**
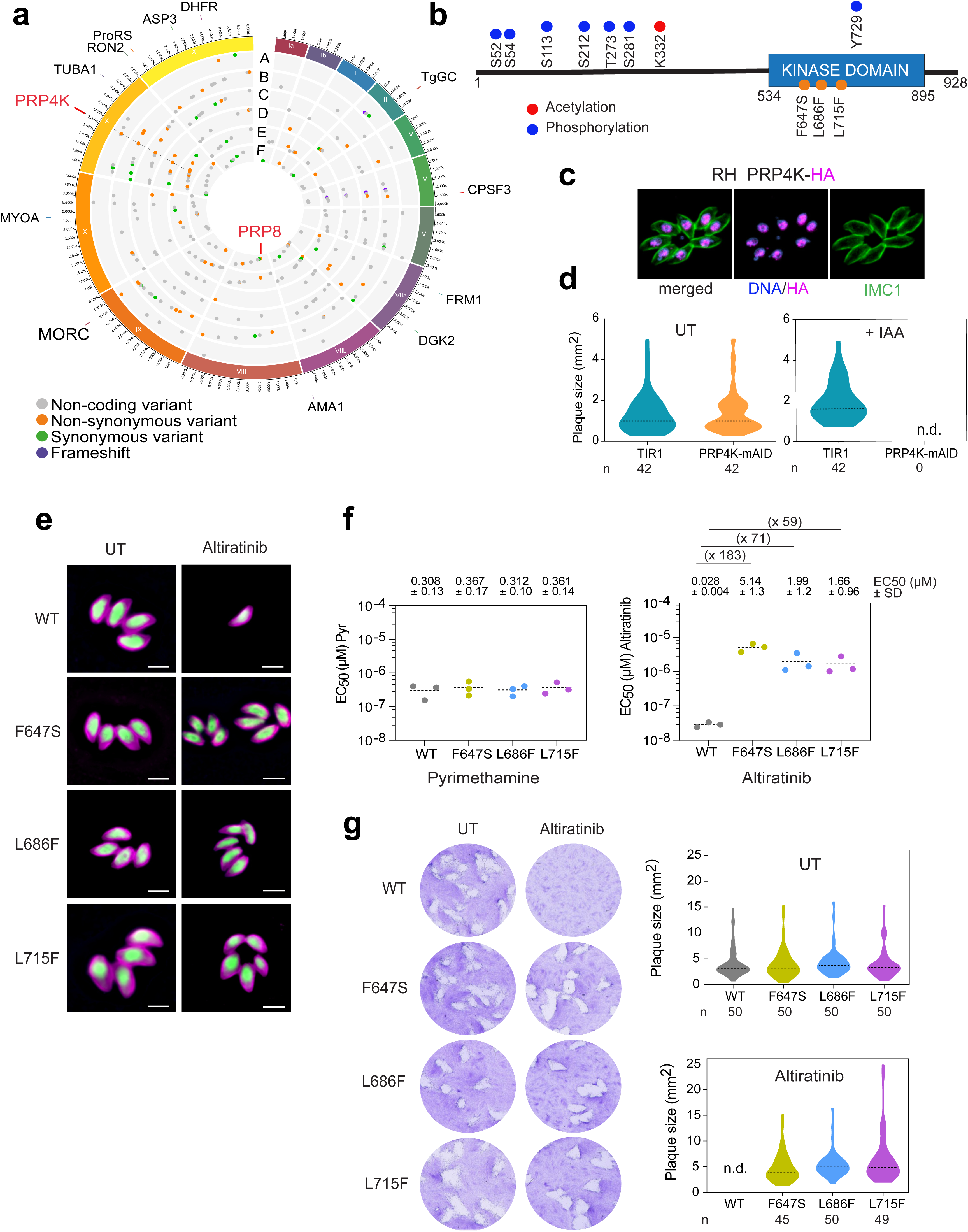
Deconvolution and validation of the *Tg*PRP4K molecular target. **a,** Circos plot summarizing the single nucleotide variants (SNVs) detected by transcriptomic analysis of *T. gondii* altiratinib -resistant lines, grouped by chromosome (numbered in Roman numerals with size intervals indicated on the outside). Each dot in the six innermost gray tracks corresponds to a scatter plot of the mutations identified in the six drug-resistant strains, with each ring representing one of the six drug-resistant lines (A through F). Each bar in the outermost track represents the positions of selected archetypal essential genes. See Supplementary Table 2 for transcriptomic analysis. **b**, Schematic representation of the *Tg*PRP4K protein structure. The kinase domain is predicted in the C-terminal portion of the protein. Phosphorylated and acetylated residues are shown as blue and red dots, respectively. The orange dots correspond to the three discovered SNVs located in the kinase domain**. c,** The nuclear location of *Tg*PRP4K (red) in human primary fibroblasts (HFFs) infected with parasites expressing an HA–Flag-tagged copy of *Tg*PRP4K. Cells were co-stained with Hoechst DNA-specific dye (blue) and the anti-Inner Membrane Complex (IMC) (green) antibody. Scale bar, 5 μm. **d,** Graphs representing the essentiality of *Tg*PRP4K protein assessed by plaque assay. RH_Tir1-Ty and *Tg*PRP4K KD parasites were either untreated or treated with IAA for 7 days and the size of 42 plaques were measured upon detection. n.d., not detected. **e,** Fluorescence microscopy showing intracellular growth of WT and the *Tg*PRP4K edited parasites (F647S, L686F, L715F). HFF cells were infected with tachyzoites of the indicated *T. gondii* strains expressing the NLuc-P2A-EmGFP reporter gene and incubated with 300 nM of altiratinib or 0.1% DMSO as control. Cells were fixed 24 h post-infection and then stained with antibodies against the *T. gondii* inner membrane complex protein GAP45 (magenta). The cytosolic GFP is shown in green. Scale bars represent 5 μm. **f,** EC_50_ values for pyrimethamine (Pyr) and altiratinib were determined for WT and the engineered *Tg*PRP4K mutant strains (F647S, L686F, L715F). The EC50 values on the upper part of the graphs represent the mean ± SD of three biological replicates. On the top of each panel, lines shown the fold change in EC_50_ relative to that of the WT parasites. Dose-response curves are shown in Extended Data Fig. 3c**. g,** Effects of *Tg*PRP4K mutations on *T. gondii* lytic cycle as determined by plaque assay. Plaque sizes (n = 50 per condition) were measured for WT and the engineered *Tg*PRP4K mutant strains (F647S, L686F, L715F) after 7 days of growth in the absence or presence of 300 mM of altiratinib. n.d., not detected. Significance was assessed by Mann-Whitney or Kruskal-Wallis tests (One-way ANOVA).

*TGGT1_313180* encodes a 928 amino acid (aa) protein that has a predicted kinase domain in its C-terminus, hereafter referred to as *Tg*PRP4K (Fig. 2b). *Tg*PRP4K is phylogenetically related to the cyclin-dependent-like kinase family (CLK) (Talevich, E. et al. 2011; Zhou Z and Fu XD, 2013) and its closest ancestor in humans is the splicing factor kinase PRP4 kinase (PRP4K or PRPF4B) and in *P. falciparum* is *Pf*CLK3 (*PF3D7_1114700*), a kinase that has been identified as a multistage cross-species antimalarial drug target (Extended Data Fig. 2a) (Agarwal, S. et al. 2011 ; Alam MM et al. 2019; Mahindra A et al. 2020). Immunofluorescence analysis of intracellular parasites showed that *Tg*PRP4K is localized to nuclear speckle-like structures (Fig. 2c). *TgPRP4K* is essential for the lytic cycle of tachyzoites, as its genetic deletion results in a fitness score of -4.69 (Sidik *et al.,* 2016), and conditional deletion of the kinase using the auxin-inducible degron system (AID) significantly impairs parasite growth (Fig. 2d) in agreement with a recent study (Lee VV et al. bioRxiv preprint).

Surprisingly, the altiratinib-resistant parasite line from mutagenesis F has a wild-type (WT) allele of *Tg*PRP4K and a mutation E1325K in *Tg*PRP8, a protein located in the catalytic core of the spliceosome that has been shown to interact with PRP4K in *Schizosaccharomyces pombe* to facilitate spliceosome activation (Bottner *et al.,* 2005; Charenton C *et al.,* 2019). This reinforces the possibility that the PRP4K-PRP8 complex is at the basis for the anti-*toxoplasma* activity of altiratinib. The specific association between *Tg*PRP4K and *Tg*PRP8 was then confirmed by FLAG affinity immunoprecipitation and mass spectrometry (MS)-based proteomic analyzes using knock-in parasite lines expressing a tagged version of each protein (Supplementary Table 3). Other partners have been identified as pre-mRNA splicing proteins constitutive of the core spliceosome, such as U2 snRNP proteins and U5 snRNP proteins, including the RNA helicase Brr2 and Snu114, which forms a pocket enclosing the catalytic RNA network of activated spliceosomes (Supplementary Fig. 1) (Bertram K *et al.,* 2017; Charenton C *et al.,* 2019). Known pre-mRNA splicing factors were also purified along with PRP4K and PRP8 (Supplementary Table 3 and Fig. 1). *Tg*PRP4K was found in a high molecular-weight complex (∼500 kDa; fractions 24–26) that withstood stringent salt conditions and partially co-eluted with the *Tg*PRP8-containing spliceosome, which migrates by size exclusion chromatography with an apparent molecular weight of ∼900 kDa (fractions 18-20) (Extended Data Fig. 2b, c).

### Mutations within *Tg*PRP4K confer resistance to altiratinib

To confirm that the mutations found in *Tg*PRP4K and *Tg*PRP8 were sufficient to confer resistance to altiratinib, we used the CRISPR/Cas9 system in conjunction with homology-directed repair, to reconstruct each of the etiological mutations into the susceptible parental *T. gondii* strain (Bellini *et al.,* 2020). Parasites were cotransfected with a vector expressing the Cas9 endonuclease and a synthetic guide RNA (sgRNA) targeting either *TgPRP4K* or *TgPRP8*, and the corresponding homologous single-stranded donor oligonucleotides (ssODN) as repair templates (Extended Data Fig. 3a). After altiratinib selection, the resistant parasites were cloned, and DNA sequencing confirmed that the mutations were properly introduced at the *TgPRP4K* locus (Extended Data Fig. 3b). Note that despite numerous attempts, allelic substitution for *TgPRP8* could not be achieved, suggesting that the *Tg*PRP8 (E1325K) mutation alone does not confer resistance to altiratinib and was not investigated further. Compared with WT parasites, mutant strains edited for *TgPRP4K* (mutations F647S, L686F, and L715F) significantly decreased sensitivity to altiratinib by 50- to 180-fold (Fig. 2e-g and Extended Data Fig. 3c), suggesting that altiratinib targets *Tg*PRP4K activity.

### Structural investigation of the mechanism of action of altiratinib on *Tg*PRP4K

To unravel the molecular mechanism of action of altiratinib inhibition, we expressed the predicted kinase domain of *Tg*PRP4K in the WT and L715F variant versions by removing the intrinsically disordered region at the N-terminus (Extended Data Fig. 4a). Both recombinant proteins were produced in satisfactory yields, with the notable difference being a higher size homogeneity of the L715F mutant (Extended Data Fig. 4b) when analyzed by size exclusion chromatography coupled to laser light scattering (SEC-MALLS). The same sample did not display a double band which can be seen in the WT (Extended Data Fig. 4c) and was also observed in Flag-purified human WT PRPF4B as a result of posttranslational modifications (Dellaire et al., 2002). The L715F mutation is peculiar because it centered on the DFG motif, which is a DLG in apicomplexan parasites and plays a central role in regulating the activation loop. Using a thermal stability assay and a thermophoresis titration assay, we could show a direct stabilizing effect (delta Tm of 11°C) and binding (Kd of 4 μM) of altiratinib with the WT *Tg*PRP4K kinase domain (Fig. 3a, b). Counterintuitively though, the L715F mutation does not decrease altiratinib binding, but instead increases binding affinity (Fig. 3a, b). This not only increases the apparent Kd value by almost a factor of 10, but also increases the stabilizing effect of the compound *in vitro* compared to WT (with a delta Tm of 15 °C). This extraordinary observation highlights an unusual resistance mechanism that compensates for the inhibitory mechanism regardless of the binding affinity of the compound. Using this point mutant, we successfully co-crystallized *Tg*PRP4K in complex with altiratinib and obtained high-resolution diffraction to 2.2Å (pdb id: 7Q4A, Supplementary Table 4). A molecular replacement solution was found with the human homolog of PRPF4B kinase domain (pdb id: 6CNH), which shares 47% sequence identity with *Tg*PRP4K. The structure solution showed *Tg*PRP4K crystallizing as a dimer with the catalysis cavities facing each other (Extended Data Fig. 5a). The monomer B exhibited more complete density within the flexible regions, so all further structural representations are based on this monomer. The activation loop was fully assembled in our model and occupies a DFG “out” conformation (Fig. 3c, Extended Data Fig. 5c) while the tyrosine 729 is phosphorylated in this structure (Fig. 3c, Extended Data Fig. 5b). Interestingly, this phospho-tyrosine is central to the ability of *Tg*PRP4K to crystallize under these conditions, as it forms numerous crystal contacts with other symmetry-related molecules (Extended Data Fig. 5b).

**Fig. 3.**
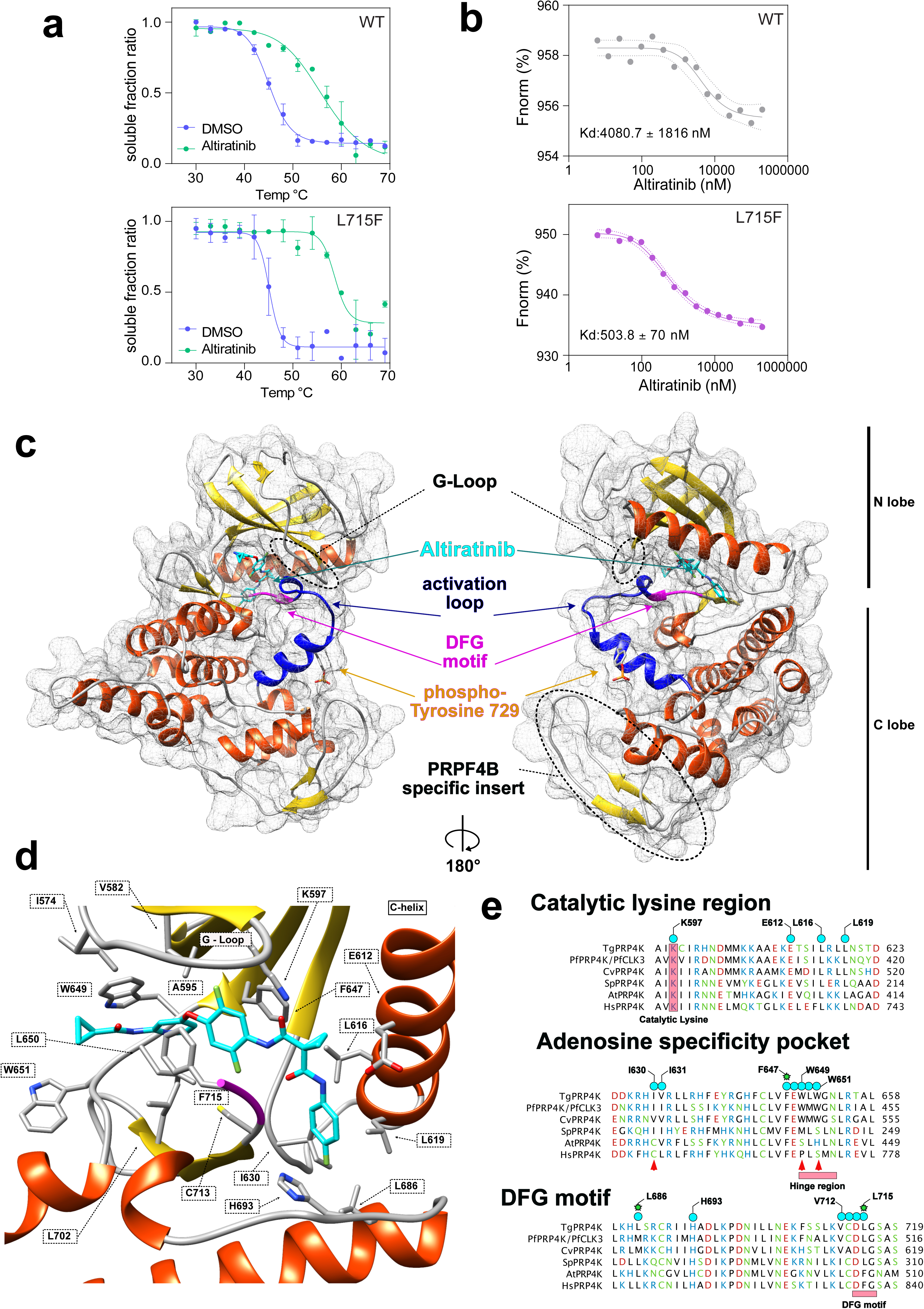
Structure of the complex *Tg*PRP4K-altiratinib and mechanism of action. **a,** Thermal stability profile of WT (upper panel) and L715F (bottom panel) recombinant proteins. Each protein was incubated for 3 minutes at different temperatures (from 30° to 69° C) in presence or absence of Altiratinib (100 µM) to quantify the melting temperatures using non-linear regression analysis of normalized data and assuming a sigmoidal dose response. **b,** Protein–Ligand interaction. WT and L715F recombinant proteins labelled to His-fluorescent dye (200nM), were incubated with altiratinib from 105 nM to 6.1 nM. Changes in thermophoresis were plotted, yielding a Kd of 4808 ± 1816 nM for WT (upper panel) and 503.8 ± 70 nM for L715F (bottom panel). Error bars: dotted lines; n= 3. **c,** Full structure of *Tg*PRP4K (L715F) bound to altiratinib (pdb id: 7Q4A). PRP4K is represented in a cartoon fashion with a transparent surface background with alpha helices colored in orange and beta strands colored in yellow. The activation loop is highlighted in blue, the DFG backbone is shown in pink, the phosphor-serine 729 side chain and altiratinib are shown in a stick representation and colored in grey and cyan respectively. **d,** Altiratinib binding within *Tg*PRP4K. Zoomed in focus on altiratinib and the key interacting side chains of *Tg*PRP4K shown as grey sticks. Cartoon colors are the same as used in panel a. **e,** Sequence alignment of altiratinib *Tg*PRP4K binding regions compared against *Plasmodium falciparum* (*Pf*), *Chromera Velia* (cc), *Schizosaccharomyces pombe* (*Sp*), *Arabidopsis thaliana* (*At*) and *Homo sapiens* (*Hs*) PRP4K/CLK3 orthologs. Key regions are highlighted by pink rectangles, altiratinib interacting amino acids from *Tg*PRP4K are shown by cyan circles while divergent residues in the human ortholog are shown by red triangles. Mutations found in the mutagenesis experiment are highlighted by a green star.

The activation loop displays an alpha helix (Ile 726 to Tyr 735) that appears to be unique to *T. gondii* PRP4K when compared to the human ortholog, which was only ever crystallized in DFG “in” conformations and is largely a random coil in this state (Fig. 3c, Extended Data Fig. 5c). Compared to the human ortholog in its global structure, *Tg*PRP4K is structurally conserved, with minor structural differences in the C-terminal portion (aa 840 to 854) of the kinase domain (Extended Data Fig. 5c). The structure also reveals a C-terminal antiparallel short beta strand that, to our knowledge, is unique to the PRP4K kinase lineage (Fig. 3c) and is also structurally conserved in the human ortholog (Extended Data Fig. 5c), although sequence conservation is very low.

Electron density for altiratinib was clearly visible in our crystal structure and interacts in the ATP-binding pocket located at the interface between the N- and C-lobes (Fig. 3c), with the DFG motif and the G-rich loop closing off this cavity. Remarkably, both monomers display strong electron density for altiratinib, allowing us to confidently assign the entire molecule (Extended Data Fig. 6a). More detailed analysis revealed that the interaction of the compound within the cavity relies on numerous hydrophobic interactions (Fig. 3d, Extended Data Fig. 6b and 6c), which can be divided into three distinct zones. The first zone, consisting of a cyclopropanecarbonylamino group connecting a pyridine ring, interacts mainly with side chains W649, L650, W651, L702 and A595 (Fig. 3d, Extended Data Fig. 6c). Hydrogen bonds also form with the carbonyl and amide groups of the leucine 650, and most of these residues form the ATP-binding hinge region leading to the deeper allosteric pocket. The second zone of altiratinib is central and consists mainly of a difluorophenyl ring stacked between the two phenylalanines 647 and 715 (the DFG central residue, which is a leucine in wild type *Tg*PRP4K) (Fig. 3d), with one of the fluorine groups interacting with the sulfur group of C713 (Extended Data Fig. 6c). These interactions ensure that the activation loop remains in this « out » position. Finally, the last part of altiratinib which encompasses a cyclopropane-1,1-dicarboxamide leading to a fluorophenyl ring, is buried deep in the allosteric cavity and interacts with multiple residues within the C-lobe, notably the glutamic acid 612 and leucine 616 and 619, which line up on the C-alpha-helix (Fig. 3d, Extended Data Fig. 6c). Other interactions are mediated by I630 and L686, as well as the H693, which belongs to the canonical HxD triad that is a H/A/D in PRP4K proteins. Only one residue within the N-lobe, the catalytic lysine K597, forms a hydrogen bond with the central carboxy group. Using this structure, we can now rationalize the consequences of the resistance mutations triggered by our EMS screen. All of the point mutations we obtained involve residues that interact directly with altiratinib, whereas the direct mechanisms of resistance are likely quite different (Fig. 3e). The L686F mutation logically introduces a steric hindrance for the fluorophenyl ring by significantly increasing the size of the side-chain. The other two resistance-conferring mutations, F647S and L715F, involve residues in direct interaction and at opposite sides of the central difluorophenyl ring. F647S probably strongly decreases hydrophobic stacking, while we have evidence that the mutation L715F does not cause steric hindrance but, on the contrary, probably increases hydrophobic stacking of the difluorophenyl ring. However, within these mutants there is little evidence pointing towards a species specificity, although altiratinib is not recognized as an inhibitor of PRPF4B in human cells, as it was originally designed to inhibit the kinases MET, TIE2 (TEK), and VEGFR2 (KDR) (Kwon Y et al. 2015; Smith DB et al. 2015). Of the residues involved in binding to altiratinib, most are strictly conserved among PRPF4B orthologs (Fig. 3e), but the hinge region has residues (W649 and W651) that diverge considerably from the human ortholog, being replaced by a proline and serine, respectively.

### Hinge region residue 649 controls species specificity of altiratinib towards *Tg*PRP4K

The superposition of the human and *T. gondii* PRPF4B/PRP4K structures makes it clear that the hinge region has a consistent backbone structure despite significant differences in side chain composition (Fig. 4a). More importantly, this overlay shows that the change from W649 to P769 would affect the main hydrophobic component that stacks the cyclopropanecarbonylamino and pyridin groups of altiratinib. A similar, albeit lesser, role can also be attributed to W651, whose equivalent residue in humans is S771 and likely reduces the hydrophobic caging potential toward altiratinib. To test the significance of residue W649, we used the same CRISPR-Cas9 complementation approach for SNP validation to generate a “humanised” mutant W649/P that requires a codon change from TGG to CCG (Fig. 4b). The probability of such a change occurring in EMS mutagenesis is low because a change from G to C is required between two replaced nucleotides. This substitution is not prevalent in EMS mutagenesis, which preferentially alkylates G residues (Greene EA et al. 2003). Remarkably, this artificial humanization produced parasites that were resistant to altiratinib (Fig. 4c-e) and had an EC50 of 3.5 μM, which is comparable to the mutations using the EMS approach. Finally, using *Tg*PRP4K WT, L715F and W649P expressed in insect cells, we were able to probe the *in vitro* consequences of these two different mutations on the ability of the protein to interact with altiratinib. Using an indirect thermal shift assay (Fig. 4f, h) and a thermophoresis approach (Fig. 4g), we demonstrated that the hydrophobic stacking of W649 is essential for altiratinib binding, as almost no stabilization is observed in the presence of altiratinib (delta Tm of 3 °C), compared to WT PRPF4B (Delta Tm of 11°C), while the binding affinity measured in thermophoresis transitions from 4 μM to not measurable.

**Fig. 4.**
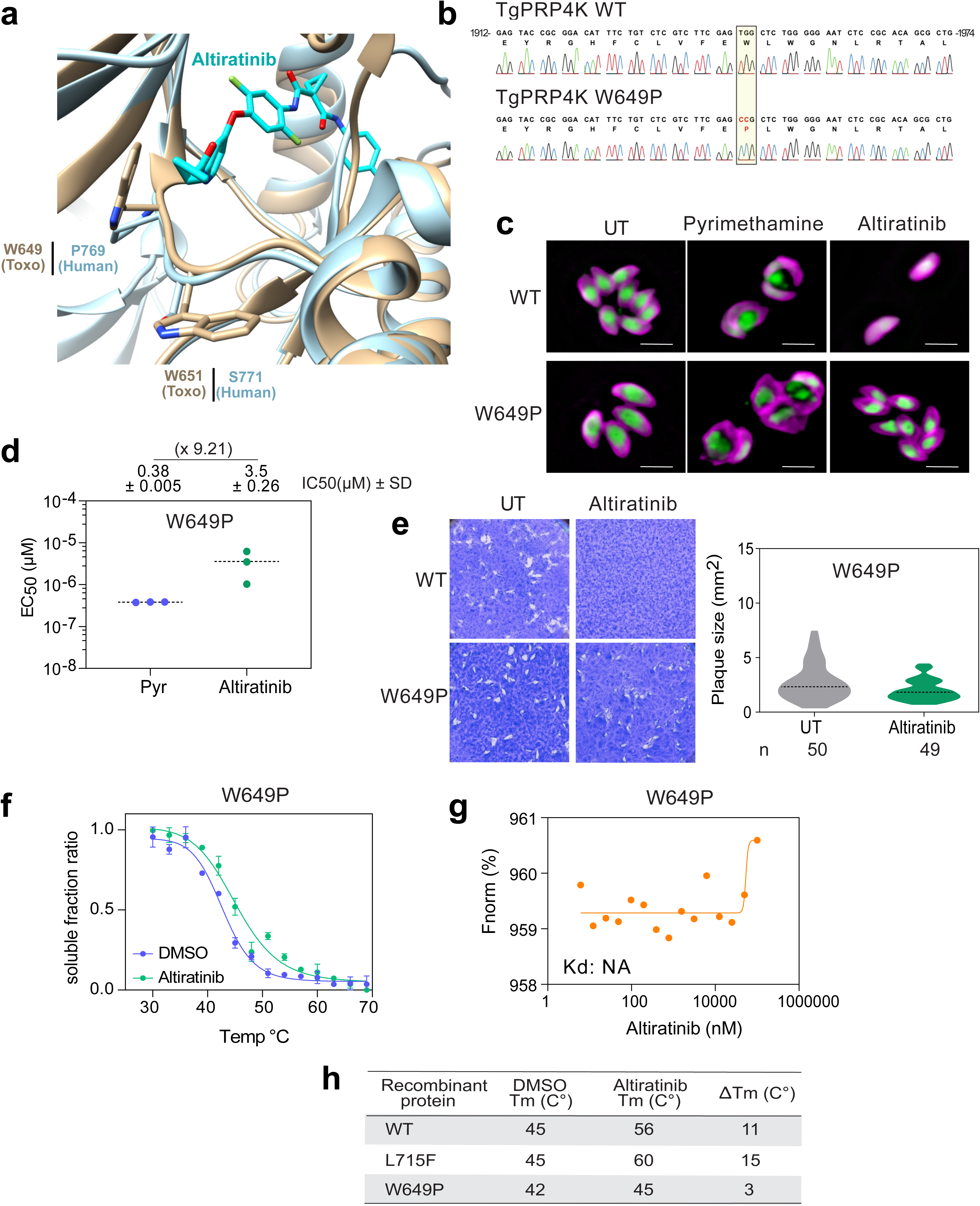
Hinge region selectivity towards altiratinib. **a,** Hinge region species selectivity towards altiratinib. Cartoon diagram of structurally superposed *Tg*PRP4K (tan) and human PRPF4B (sky blue) with altiratinib in cyan. Hinge region residues are detailed by including stick representations of their side chains. **b,** Sanger chromatogram validating the *TgPRP4K* gene editing for W649P mutation. On the top, nucleotide positions relative to the ATG start codon on genomic DNA are indicated. **c,** IFA showing the W649P resistance to altiratinib. Confluent HFFs were infected with engineered parasites and incubated with pyrimethamine (1 µM) or altiratinib (300 nM) for 24h. Fixed cells were stained using anti-GAP45 antibody (magenta) while the cytosolic GFP is showed in green. Scale bar represents 5 µm. **d,** Graph representing the EC_50_ of W649P for pyrimethamine and altiratinib. Values showed in the upper part on the graph are the mean ± SD of three independent experiment. On the top of the panel, the line shows the fold change in altiratinib EC_50_ relative to pyrimethamine. **e,** Plaque assay representing the lytic cycle of RH WT and W649P parasites in presence or absence of altiratinib. After 7days of drugs incubation, infected cells were fixed and stained to visualize the presence of lysis plaques (on the left). The area of 50 plaques was measured and represented in the right panel. **f,** Thermal stability profile of W649P recombinant protein in presence or absence of Altiratinib. **g,** Protein-ligand interaction profile of W649P protein in presence of Altiratinib. Changes in thermophoresis of three replicates were plotted. Error bars: dotted lines. NA, not available. **h,** Table showing the melting temperature (Tm) of WT, L715F and W649P recombinant proteins during their incubation with DMSO or Altiratinib at different temperatures. Low interaction between W649P and the compound was detected as showed by the ΔTm values.

### Chemical inactivation of *Tg*PRP4K activity disturbs pre-mRNA splicing in *T. gondii*

Since it has been proposed that the human kinase PRPF4B and *Pf*CLK3 regulate RNA splicing (Schneider M *et al*. 2010; Alam *et al*. 2019), we examined transcriptional changes in the parental parasite RH and in the drug-resistant strains L715F and W649P in response to exposure to altiratinib using nanopore long-read direct RNA sequencing (DRS), a technology well suited for determining the full repertoire of mRNA species, including alternative splicing isoforms and divergent patterns, if present. The most obvious effect was that a substantial number of genes (n=2400) showed altered mRNA expression, of which 784 were induced and 1616 suppressed when the parent strain was treated with altiratinib, whereas no change was observed in the two mutant strains exposed to the drug (Fig. 5a). This confirms that altiratinib disrupts mRNA transcription, which was expected, but also that the drug specifically targets *Tg*PRP4K, as both mutations not only confer resistance (Fig. 2d-f) but also restore gene expression to the untreated state (Fig. 5a). Having identified isoforms with high confidence using the Nanopore data, we used FLAIR (Full-Length Alternative Isoform Analysis of RNA) (Tang AD *et al.,* 2020) as a framework for analyzing differential isoform usage in wild-type and mutant strains left untreated or exposed to altiratinib. The most important transcriptional phenotype was the change in pre-mRNA splicing dynamics associated with inhibition of *Tg*PRP4K exclusively in WT parasites (Fig. 5b). At many *loci*, chemical inactivation of *Tg*PRP4K was accompanied by complete retention of the second intron (e.g., *TGME49_214940*; Fig. 5b,d) or intron retention and exon skipping at the same *loci* (e.g., *TGME49_211420* and *TGME49_247350*; Fig. 5c; Extended Data Fig. 7a). When an intron is spliced, it rapidly promotes splicing of subsequent introns, whereas when splicing is hindered, subsequent introns tend to be retained, leading to the concept of ‘all or none’ splicing (Oesterreich FC et al. 2016; Herzel L et al. 2018). Consistent with this concept, we regularly observed a global collapse of splicing along the entire transcript (e.g., *TGME49_208450*; Extended Data Fig. 7b) after drug treatment.

**Fig. 5.**
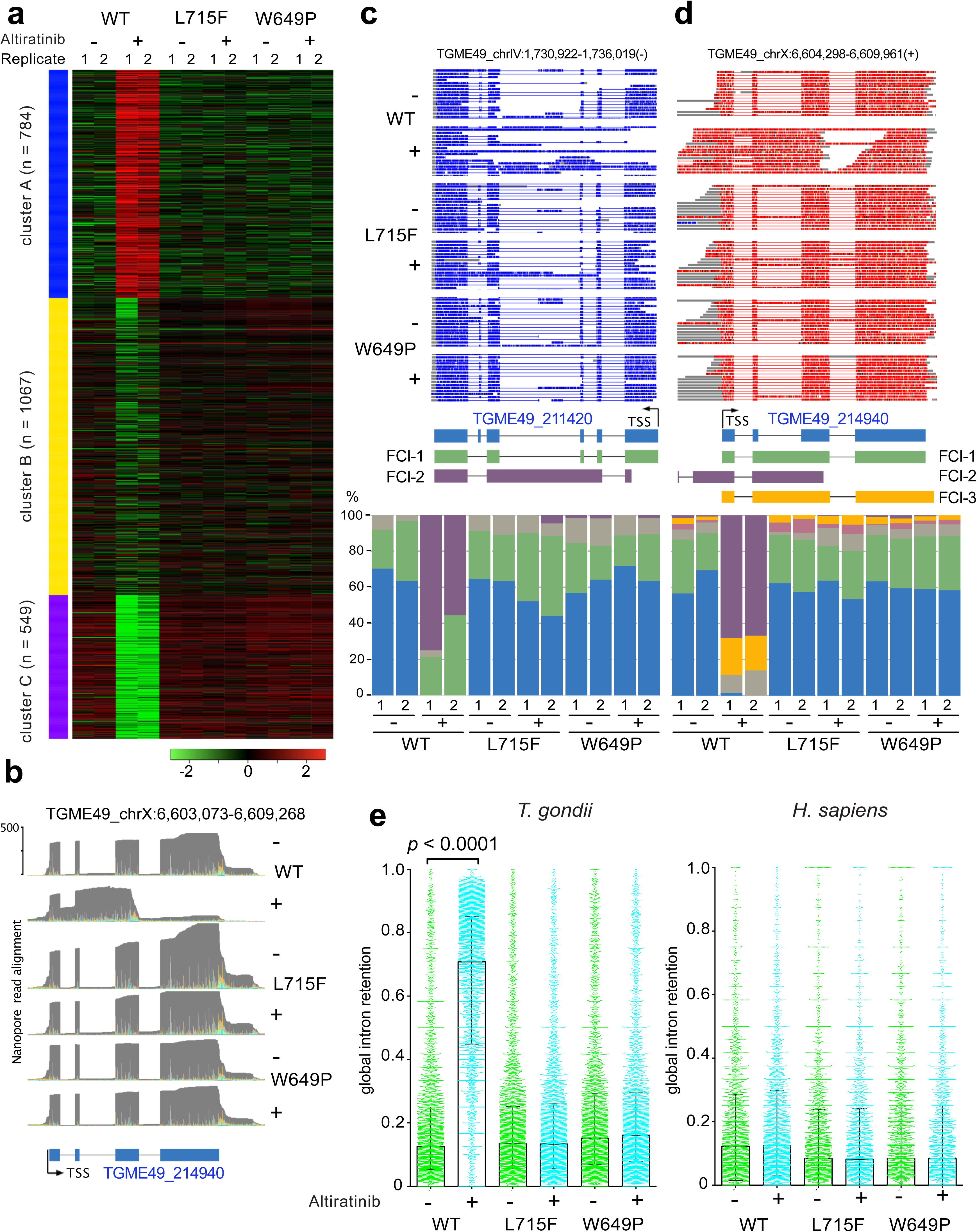
Nanopore DRS analysis of altiratinib-induced splicing defects in *T. gondii*. **a,** General transcriptomic effects of altiratinib treatment. *k*-means clustering of 2400 transcripts treated with EdgeR: log2(CPM+4). The color key ranges from -3 to 3 (green to red), 3 clusters were defined. In each, *Tg*PRP4K WT/L715F/W649P duplicate sequencing experiments are shown in the presence or absence of altiratinib. **b,** M-pileup representation of the aligned nanopore reads at the *TGME49_214940* loci. WT/L715F/W649P sequencing experiments are shown as grayscale histograms in the presence or absence of altiratinib. **c and d,** FLAIR analysis of *TGME49_211420* (**c**) and TGME49_214940 (**d**) loci. Standard annotation and FLAIR collapsed isoforms (FCI) are shown schematically under a sample view of 15 reads per condition (same conditions as in **b**.). Sense and antisense reads are colored red and blue, respectively. Below the FCI representation is an isoform quantification histogram showing duplicate measurements in each WT/L715F/W649P and untreated (-) or treated (+) condition. The color code is the same as for the above FCI, grey histograms represent minor isoforms which not shown schematically. **d.** Overall quantification of intron retention. Scatter plot of intron retention ratios (per averaged duplicate transcript) are shown for *T. gondii* and *H. sapiens*. WT/L715F/W649P strains that were untreated (in green) or treated (in cyan) are shown, the black histogram shows the median, the whiskers show the interquartile range. Significance between the WT untreated and treated conditions was calculated using a non-parametric Friedman test.

Since splicing is predominantly cotranscriptional, we also observed that intron retention leads to premature transcriptional termination (e.g., *TGME49_278940*; Extended Data Fig. 7c). At the transcriptome level, intron retention is the predominant aberrant splicing event found in altiratinib-treated WT tachyzoites in contrast to the host cells they infect, underscoring the high degree of selectivity of altiratinib (Fig. 5e). Upon closer inspection, we found that intron retention leads to premature termination of translation due to frameshifts, which may ultimately lead to altered function of the protein-coding gene. In addition, aberrant isoforms are degraded, as indicated by the lower read rates at some *loci*, likely through nonsense-mediated decay (NMD), a quality control mechanism that eliminates transcripts with a premature termination codon. In this way, treatment with altiratinib leads to the production of defective proteins that ultimately affect parasite survival.

### Altiratinib also causes mis-splicing in *P. falciparum* but not in *C. parvum*, which has a divergent PRP4K ortholog

Because altiratinib was active against a wide range of apicomplexans (Fig. 1) and the PRPK4/CLK3 family was well conserved within the phylum, we wondered whether the drug might inhibit splicing in other parasites of the phylum. We first examined transcriptional changes of red blood cells infected with *P. falciparum* after treatment with altiratinib using Nanopore DRS. All types of splicing defects that we had observed in *T. gondii* were also present in *P. falciparum*, such as exon skipping, intron retention, and premature transcription termination (Fig. 6a; Extended Data Fig. 8a-c), with a general trend toward global splicing collapse along the entire transcript, with premature mRNAs being highly susceptible to NMD degradation (Extended Data Fig. 8a-c). As with *T. gondii*, markedly increased intron retention is a conserved phenomenon in *P. falciparum* exposed to altiratinib (Fig. 6b). These results underscore the potential targeting by altiratinib of *Pf*CLK3 (*PF3D7_1114700*), a kinase that is essential for *P. falciparum* survival in red blood cells and plays a critical role in regulating RNA splicing of the malaria parasite (Alam MM, 2019; Mahindra *et al.,* 2020). We then took the opportunity to test the drug on *Cryptosporidium parvum*, a parasite of the phylum that differs from others in having a significantly divergent ortholog of PRP4K/CLK3 (*cgd8_5180*), specifically the resistance-conferring DFG motif instead of the DLG motif found in *T. gondii* and *P. falciparum,* but also several significant mutations at other altiratinib-interacting residues (L719 to F, W651 to H, and C713 to S) that may strongly affect the binding selectivity of altiratinib (Extended Data Fig. 2a and Extended Data Fig. 8d). As expected, we observed no defects in mRNA splicing in *C. parvum* exposed to altiratinib (Fig. 6c-d; Extended Data Fig. 8e), again confirming the selectivity of the drug for PRP4K/CLK3 with a DLG motif and ruling out off-target activities.

**Fig. 6.**
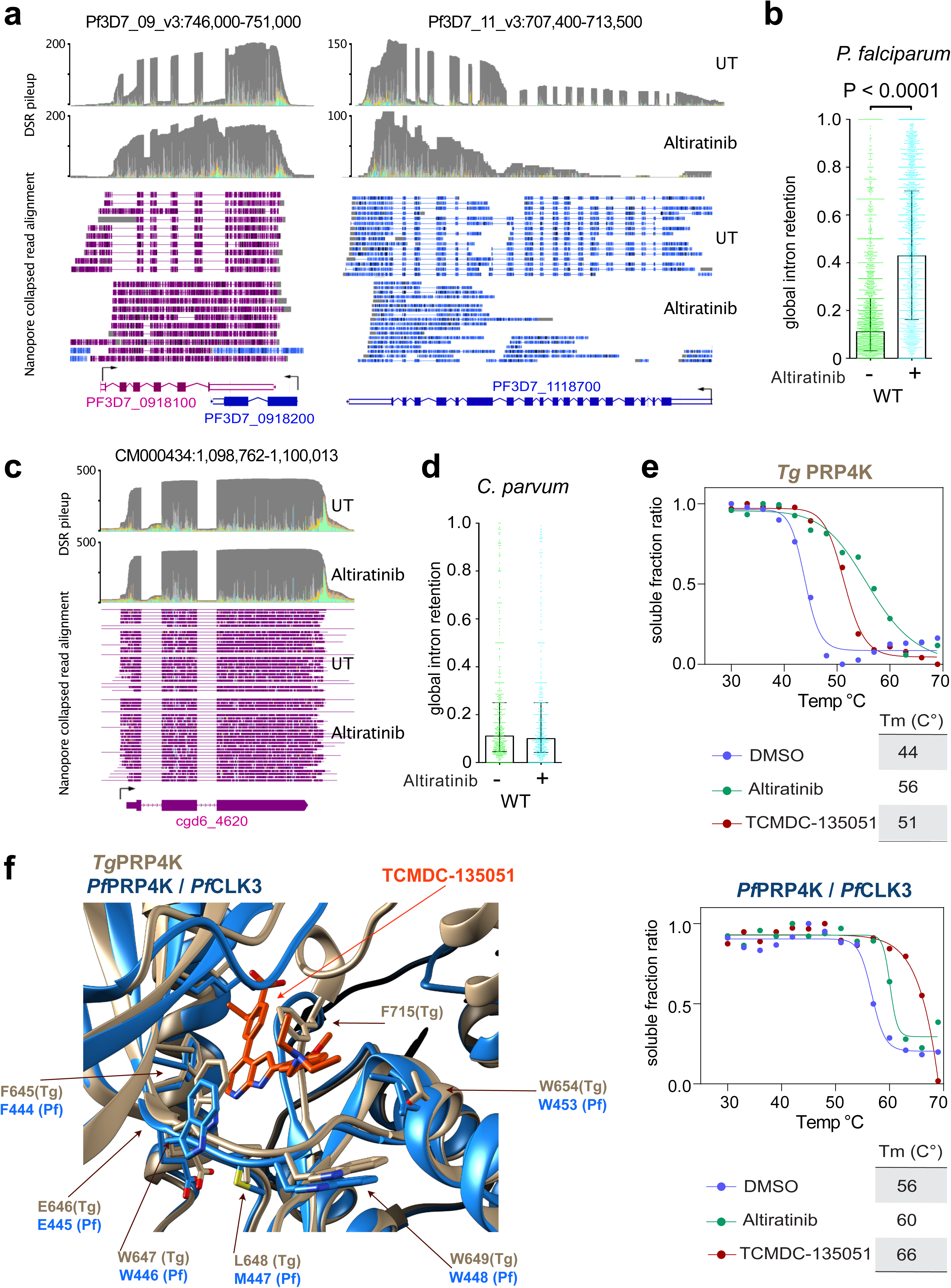
Cross-species selectivity of altiratinib analysed by nanopore DRS. **a,** Splicing defects induced by altiratinib in *P. falciparum*. M-pileup representation of aligned nanopore reads at the *PF3D7_0918100* and *PF3D7_1118700* loci. Untreated (UT) or altiratinib-treated sequencing experiments are shown as grayscale histograms. Shown below are IGB samples from 10 individual aligned reads using sense (purple) and antisense (blue) coloring under UT and treated conditions. **b,** Overall quantification of intron retention in *P. falciparum*. Scatter plots of intron retention ratios (per averaged duplicate transcript) are shown for untreated (in green) or treated (in cyan) conditions, the black histogram shows the median, the whiskers show the interquartile range. Significance between the WT untreated and treated conditions was calculated using a nonparametric Mann-Whitney *t*-test. **c,** Splicing consistency is maintained in *C. parvum*. M-pileup and IGB sampling of aligned reads from untreated (UT) or treated *C. parvum* at the highly transcribed and spliced *cdg6_4620* loci. **d,** Overall quantification of intron retention in *C. parvum*. The same display rules as in **b**. were applied. **e.** Hinge region species selectivity towards TCMD-135051. Cartoon diagram of the structurally superposed *Tg*PRP4K from this work (in tan) and the alphaflod2 predicted *Pf*CLK3 (dodger blue) with TCDM-135051 modelling in orange. The residues of the hinge region are also detailed by showing their side chains as stick representations. **f,** Thermal shift assay of *Tg*PRP4K and *Pf*CLK3 in the presence of altiratinib or TCDM-135051.

### The ins and outs of CLK3/PRP4K inhibition by altiratinib or TCMDC-135051

To further confirm *P. falciparum Pf*CLK3/PRP4K as a target of altiratinib, we expressed the WT *Pf*PRP4K kinase domain to probe this biochemical interaction (Extended Data Fig. 9). Using the previously described thermal shift assay, we found that altiratinib indeed stabilizes *Pf*CLK3, albeit with a weaker potential, the delta Tm is of 4°C, compared to *Tg*PRP4K, which has a delta Tm of 11°C (Fig. 6e). Interestingly, however, when probing the TCMDC-135051 compound, a recently discovered inhibitor of PfCLK3 (Mahindra *et al*., 2021), we observed a reversed trend with a stronger stabilizing effect on *Pf*CLK3 with a delta Tm of 10°C instead of 7°C for *Tg*PRP4K (Fig. 6e). These results highlight two important aspects. First, we confirm that TCMDC-135051 likely binds the active site of *Pf*CLK3 as the energy requirements for such a stabilizing effect would probably only occur within a buried cavity strongly interacting with the compound. Second, this also highlights that there may still be some species selectivity between the two compounds. As we were unable to crystallize *Pf*CLK3 in the bound or unbound state, we used alphafold2 (Jumper J *et al.,* 2021) within collabfold (Mirdita *et al.,* 2021) to create a model that we superposed to our crystallographic structure and manually docked TCMDC-135051, taking advantage of the structural homology to other hinge regions binders containing a 7-azaindole scaffold (as initially proposed in Mahindra *et al*., 2021). In this modeling (Fig. 6f), we observe that most of the PRP4K/CLK3 hinge region is conserved between *P. falciparum* and *T. gondii*, in particular residues W649/W647 in *Tg*PRP4K, which are also fully conserved in *Pf*CLK3 (W446/W448) and likely also have an important impact on the selectivity of TCMDC-135051, particularly through hydrophobic stacking. However, the conformation of the activation loop is not consistent with the binding of TCMDC-135051 in the *Tg*PRP4K structure (Fig. 6f), indicating potential differences in the activation loop conformation that may differ between TCMDC-135051 and altiratinib.

## Discussion

Our studies define altiratinib as a promising pan-apicomplexan drug candidate effective against the human pathogens *T. gondii* and *P. falciparum*, as well as *N. caninum* and *E. tenella* of veterinary interest. Using genetic, structural and transcriptional approaches, we have shown that repurposing of altiratinib disrupts mRNA splicing in *T. gondii* and *P. falciparum* by targeting the kinase core of PRP4K/CLK3. The induced splicing defects are so extensive that they lead to irreversible inhibition in the nanomolar range of rapidly proliferating apicomplexan zoites in cellular assays. Using a genetic target-deconvolution strategy, we have highlighted key residues involved in binding to altiratinib. Unexpectedly, this has allowed us to crystallize and resolve the first structure of a previously elusive apicomplexan kinase PRP4K/CLK3. This co-crystal structure allows us to assign an electron density to altiratinib located at the interface between the N and C lobes and occupying both the ATP-binding site and the allosteric pocket, a singular type of binding that holds PRP4K in a DFG-out conformation consistent with inhibition of the type II kinases. The structural data have clarified many unanswered questions related to the species selectivity of altiratinib, as we now know that its ability to discriminate the human ortholog and bind the parasitic PRP4K/CLK3 is constrained by residues W649/W651 in the hinge region (Fig. 4a), which have diverged significantly and are also likely critical for binding of the recently discovered *Pf*CLK3 inhibitor TCMDC-135051 (Fig. 6f) (Alam MM, 2019; Mahindra *et al.,* 2020). Another important divergence is the shift from DFG to DLG that has occurred between mammals and some apicomplexans. DLG is indeed associated with inactive or weakly active kinases such as ROR2 (Artim SC et al. 2012; Mendrola JM, 2013), and the selective pressure that led to this mutation is not yet clear, as DFG-mutated tachyzoites behave normally in cell cultures (Fig. 2d-f). This likely gain in activity is sufficient to resist altiratinib, although binding affinities are increased.

Some open questions remain to be answered. Although the evidence for a direct interaction between altiratinib and PRP4K/CLK3 and drug-induced mis-splicing is overwhelming, the true mechanism of spliceosome inhibition is still in question, as PRP4K not only plays a role in pre-B spliceosome activation by phosphorylating other components of PRP, notably PRP6 and PRP31 (Schneider et al. 2010), but also structurally integrates the complex (Charenton C *et al.,* 2019) and contacts the RNase PRP8, which may be allosterically involved in its activity. Inhibition of activity or conformational entrapment (or both) may therefore be the key to proper inhibition. *Pf*CLK3 has been identified as a multistage cross-species malarial drug target and TCMDC-135051 a drug candidate with a high curative and transmission-blocking potential (Alam MM, 2019; Mahindra *et al.,* 2020). Altiratinib and TCMDC-135051 have a very different chemical space and although they rely on comparable elements within the hinge region to selectively bind apicomplexan PRP4K/CLK3, species selectivity is still present, possibly due to differences in the dynamics of the activation loop with which binding is compatible. Dual (SAR)-directed optimization will therefore open the possibility of developing a pan-apicomplexan therapy based on the altiratinib/TCMDC-135051 combination.

Our work highlights the utility of drug repurposing and provides structural mechanistic insights into understanding how the PRP4K/CLK3 family is susceptible to selective pharmacological inhibition by small drug-like molecules. This opens new opportunities to chemically improve existing molecules to optimize pathogen killing via the PRP4K/CLK3 pathway.

## Methods

### Toxoplasma gondii, Plasmodium falciparum and human cell culture

Human primary fibroblasts (HFFs, ATCC® CCL-171™) were cultured in Dulbecco’s Modified Eagle Medium (DMEM) (Invitrogen) supplemented with 10% heat inactivated Fetal Bovine Serum (FBS) (Invitrogen), 10 mM (4-(2-hydroxyethyl)-1-piperazine ethanesulphonic acid) (HEPES) buffer pH 7.2, 2 mM L-glutamine and 50 μg/ml of penicillin and streptomycin (Invitrogen). Cells were incubated at 37°C in 5% CO2. The *Toxoplasma* strains used in this study and listed in Supplementary Table 5 were maintained *in vitro* by serial passage on monolayers of HFFs. The cultures were free of mycoplasma, as determined by qualitative PCR. *P. falciparum* parasites were cultured using standard culture conditions. The drug sensitive laboratory strain 3D7 was used in this study.

### Growth of *Cryptosporidium parvum* and RNA preparation

Hct-8 cells were grown in a T- 75 flask (90% confluence) and infected with the *C. parvum* INRAE strain at a ratio of 5 oocysts per cell. Altiratinib was added to the treated flask at a concentration of 500 nM concentration. Four hours later, the flasks were washed to remove the oocysts and further incubated in presence of altiratinib until 11h post infection. All cells were collected and RNA extracted in Trizol solution for nanopore analyses.

### In vitro inhibition of Eimeria tenella

Clec213 chicken epithelial cells were grown to sub-confluence in P96-well plates and infected with *Eimeria tenella* sporozoites (INRAE strain) expressing luciferase at a ratio of 1 sporozoite per cell. Four hours later, the plates were washed and further incubated with different concentrations of altiratinib 44 hours after infection, when luciferase activity was determined to quantify parasite development (six specimens were used for each drug concentration).

### Medium-throughput screening

The TargetMol (Boston, MA) compound library consists of 514 compounds (each as a 1 mM stock solution in DMSO). Primary screening was performed in white 96-well plates (3610, Corning ® *Costar* ®). The confluent HFFs monolayer was infected with 2000 RH NanoLuc parasites strain for 2 hours before the compounds were added at a final concentration of 5 µM in a final volume of 100 µl. The culture was incubated at 37°C for 48h. The medium was removed to add 50µl of PBS and measure the growth of the parasite using the Nano-Glo® Luciferase Assay System, according to the manufacturer’s instructions (Promega). Lysis was performed in the wells by adding 50 μL of Nano-Glo® Luciferase Assay Reagent with 1:50 dilution of Nano-Glo® Luciferase Assay Substrate. After 3 minutes of incubation, luminescence was measured using the CLARIOstar® (BMG Labtech) plate reader. The bioluminescence values of the uninfected host cells were used to determine the background signal.

### Measurement of EC_50_ for *Toxoplasma gondii* parasites

To measure the EC_50_ of *Toxoplasma gondii* parasites, confluent HFFs monolayer was infected with 2000 tachyzoites of RH parasites expressing the NLuc luciferase (RH NLuc) for 2h. After parasite invasion, each compound was added to the culture in exponential concentrations. After 48h incubation at 37°C, the medium was replaced with 50 µl of PBS. The reading of luminescence was performed using the Nano-Glo® Luciferase Assay System according to the manufacturer’s instructions (Promega). After 3 minutes of incubation, luminescence was measured using the CLARIOstar® (BMG Labtech) plate reader. EC_50_ values were determined using a non-linear regression analysis of normalized data and assuming a sigmoidal dose response. EC_50_ values for each compound represent the average of three independent biological replicates. Statistical analyses were performed using the one-way test ANOVA and GraphPad 8 software.

### Measurement of CC50 for mammalian cells and determination of Selectivity Index

Human HFFs, ARPE-19, MCF7, MDA-231 and U937 cell lines (Supplementary Table 5) were plated for 1 hour in 96 well plates for 1h and incubated with exponential concentrations of the indicated compounds in a final volume of 100 µl. After 72h of culture, CellTiter-Blue Reagent® (Promega) (20 μl/well) was added directly to each well. Plates were then incubated at 37°C for 2 hours to allow cells to convert resazurin to resorufin before reading fluorescence (560(20) Ex/ 590(10) Em) with the CLARIOstar® (BMG Labtech) plate reader. The cytotoxicity concentration (CC50) of human cells was determined using nonlinear regression curve of the normalized data. CC_50_ values represent the average of two biological experiments. The Selectivity Index (SI) was determined by the average of the human CC_50_ divided by the average of the *T. gondii* EC_50_. Mitochondrial toxicity assay was performed using the “Mitochondrial ToxGlo^TM^ Assay” kit according to the manufacturer’s instructions (Promega). Briefly, 10,000 human cell lines were plated in 96 well plates with DMEM serum and glucose free, supplemented with galactose (10 mM). After 3h of culture to allow cell adhesion, increasing concentrations of altiratinib (tested drug) and sodium azide (positive control for mitochondrial toxicity) or 800 µg/ml of digitonin (positive control for cell toxicity) were added. The cell culture was maintained at 37°C for 90 min to detect cell viability (cytotoxicity) by fluorescence and ATP production by luminescence using the CLARIOstar® (BMG Labtech) plate reader.

### Reagents

The following primary antibodies were used in the immunofluorescence and immunoblotting: rabbit anti-TgGAP45 (gift from Pr. Dominique Soldati, University of Geneva), mouse anti-HA tag (Roche, RRID: AB_2314622), and rabbit anti-HA Tag (Cell Signaling Technology, RRID: AB_1549585). Immunofluorescence secondary antibodies were coupled with Alexa Fluor 488 or Alexa Fluor 594 (Thermo Fisher Scientific). Secondary antibodies used in Western blotting were conjugated to alkaline phosphatase (Promega) or horseradish peroxidase.

### Immunofluorescence microscopy

*T. gondii* infecting HFF cells grown on coverslips were fixed in 3% formaldehyde for 20 min at room temperature, permeabilized with 0.1% (v/v) Triton X-100 for 15 min and blocked in Phosphate buffered saline (PBS) containing 3% (w/v) BSA. The cells were then incubated for 1 hour with primary antibodies followed by the addition of secondary antibodies conjugated to Alexa Fluor 488 or 594 (Molecular Probes). Nuclei were stained for 10 min at room temperature with Hoechst 33258 at 2 μg/ mL in PBS. After four washes in PBS, coverslips were mounted on a glass slide with Mowiol mounting medium, and images were acquired with a fluorescence ZEISS ApoTome.2 microscope and images were processed by ZEN software (Carl Zeiss, Inc.).

### Auxin induced degradation

Depletion of *Tg*PRP4K-AID-HA was achieved with 3- Indoleacetic acid (IAA, Sigma-Aldrich # 45533) as described by Farhat *et al.,* 2020. A stock of 500 mM IAA dissolved in 100% EtOH at 1:1,000 was used to deplete mAID-tagged proteins at a final concentration of 500 μM. Mock treatment consisted of an equivalent volume of 100% EtOH at a final concentration of 0.0789%, wt/vol. To monitor the degradation of AID-tagged proteins, parasites grown in HFF monolayers were treated with auxin, or ethanol alone, for different time intervals at 37 °C. After treatment, parasites were harvested and analyzed by immunofluorescence or Western blotting. Complete elimination of *Tg*PRP4K parasites was successful after 10 hours of IAA treatment.

### Plaque assays

Confluent HFFs were infected with freshly egressed tachyzoites before adding 0.1% DMSO or the indicated compounds. Cultures were grown at 37°C for 7 days, fixed, and stained with Coomassie blue staining solution (0.1% Coomassie R-250 in 40% ethanol and 10% acetic acid). For cytotoxicity assay, the parasites were incubated with different drugs or DMSO for 16 hours. After washing the cells, the cultures were left at 37°C for 3, 6 or 10 days before fixation and staining. The size of the plaques when present was measured using ZEN 2 lite software (Carl Zeiss, Inc.) and plotted using GraphPad Prism 8.

### Toxoplasma gondii genome editing

Targeted genome modifications were performed using the *T. gondii* adapted CRISPR/Cas9 system as previously described (Farhat *et al.,* 2020). Recombinant parasites harboring allelic replacement for PRP4K^F647S^, PRP4K^L686F^, PRP4K^L715F^, and PRP4K^W649P^ were generated by electroporation of the *T. gondii* RH NLuc strain with pTOXO_Cas9CRISPR vectors targeting the *PRP4K* coding sequence (sgPRP4K^F647S^, sgPRP4K^L686F^, sgPRP4K^L715F^) and their respective donor single-stranded oligo DNA nucleotides (ssODNs) carrying respective nucleotide substitutions (PRP4K^F647S^_donor, PRP4K^L686F^_donor, PRP4K^L715F^_donor; Supplementary Table 5) for homology-directed repair. Recombinant parasites were selected with 300nM altiratinib prior to subcloning by limited dilution, and allelic replacement was verified by sequencing of *T. gondii TgPRP4K* genomic DNA.

### *Toxoplasma gondii* random mutagenesis

Parasites were chemically mutagenized as previously described (Bellini *et al*., 2020), with the following modifications. Briefly, ∼10^7^ tachyzoites growing intracellularly in HFF cells in a T25 flask were incubated for 4 h at 37°C in 0.1% FBS DMEM growth medium containing either 2.5 mM ethyl methanesulphonate (EMS) at final concentration or the appropriate vehicle controls. After exposure to the mutagen, parasites were washed three times with PBS, and the mutagenized population was allowed to recover in a fresh T25 flask containing an HFF monolayer in the absence of drug for 3–5 days.

The released tachyzoites were then inoculated into fresh cell monolayers in medium containing 300 nM of altiratinib and incubated until viable extracellular tachyzoites emerged 8–10 days later. Surviving parasites were passaged once more under continued altiratinib treatment and cloned by limiting dilution. The cloned mutants were each isolated from 6 independent mutagenesis experiments. Thus, each flask contained unique SNV pools.

### RNA-seq, sequence alignment, and variant calling

For each biological assay, a T175 flask containing a confluent monolayer of HFF was infected with RH wild-type or Altiratinib-resistant strains. Total RNAs were extracted and purified using TRIzol (Invitrogen, Carlsbad, CA, USA) and RNeasy Plus Mini Kit (Qiagen). RNA quantity and quality were measured by NanoDrop 2000 (Thermo Scientific). RNA-sequencing was performed as previously described (Bellini *et al*., 2020), following standard Illumina protocols, by GENEWIZ (South Plainfield, NJ, USA). Briefly, the RNA quality was checked with the TapeStation System (Agilent Technologies, Palo Alto, California, USA), and Illumina TruSEQ RNA library prep and sequencing reagents were used following the manufacturer’s recommendations using polyA-selected transcripts (Illumina, San Diego, CA, USA). The samples were paired-end multiplex sequenced (2 x 150 bp) on the Illumina Hiseq 2500 platform and generated at least 40 million reads for each sample. The quality of the raw sequencing reads was assessed using FastQC (http://www.bioinformatics.babraham.ac.uk/projects/fastqc/) and MultiQC (Ewels et al., 2016). The RNA-Seq reads (FASTQ) were processed and analyzed using the Lasergene Genomics Suite version 17 (DNASTAR, Madison, WI, USA) using default parameters. The paired-end reads were uploaded onto the SeqMan NGen (version 17, DNASTAR. Madison, WI, USA) platform for reference-based assembly and variant calling using the *Toxoplasma* Type I GT1 strain (ToxoDB-46, GT1 genome) as reference template. The ArrayStar module (version 17, DNASTAR. Madison, WI, USA) was used for variant detection and statistical analysis of uniquely mapped paired-end reads using the default parameters. Variant calls were filtered to select variants present in coding regions with the following criteria: variant depth ≥ 30, Q call ≥ 60, and absent in the parental wild-type strain. Mutations were plotted on a Circos plot using Circa (OMGenomics.com). For the expression data quantification and normalization, the FASTQ reads were aligned in parallel to the ToxoDB-46 build of the *Toxoplasma gondii* GT1 genome (ToxoDB-46) using Subread version 2.0.1 (Liao et al., 2013) with the following options ‘subread-align -d 50 -D 600 --sortReadsByCoordinates’. Read counts for each gene were calculated using featureCounts from the Subread package. Differential expression analysis was conducted using DESeq2 and default settings within the iDEP.92 web interface. Transcripts were quantified and normalized using TPMCalculator.

### Plasmid construction

The plasmids and primers for gene of interest (GOI) used in this work are listed in Supplementary Table 5. To construct the vector pLIC-GOI-HAFlag, the coding sequence of GOI was amplified using primers LIC-GOI-Fwd and LIC-GOI-Rev using *T. gondii* genomic DNA as template. The resulting PCR product was cloned into the pLIC-HF-dhfr or pLIC-mCherry-dhfr vectors using the Ligation Independent Cloning (LIC) cloning method. Twenty mers-oligonucleotides corresponding to specific GOI were cloned using Golden Gate strategy. Briefly, primers GOI-gRNA-Fwd and GOI-gRNA-Rev containing the sgRNA targeting GOI genomic sequence were phosphorylated, annealed and ligated into the pTOXO_Cas9-CRISPR plasmid linearized with BsaI, leading to pTOXO_Cas9-CRISPR::sgGOI. Just two transgenic components are needed to implement the auxin-inducible degron (AID) system, a plant auxin receptor called transport inhibitor response 1 (TIR1) and a POI tagged with an AID. We engineered a type I RHΔku80 and a type II lines of *T. gondii* to stably express Tir1 from Oryza sativa tagged with Ty and controlled by a promoter selected for a moderate expression of the chimeric protein. The plasmid *pModProm1-TiR1-TY1-3DHFR* (DNA sequence in Supplementary Table 5 was DNA synthetized and then cloned in pUC57 simple by Genscript. The chimeric construct was inserted within the UPRT locus. We also created a pLIC vector containing a codon-optimized for *T. gondii* expression DNA block with the mAID from *Arabidopsis thaliana* auxin-responsive protein IAA17^E66-S133^, as defined in Farhat *et al.,* 2020, in frame with a HA tag (Supplementary Table 5).

### Toxoplasma gondii transfection

*T. gondii* strains were electroporated with vectors in cytomix buffer (120 mM KCl, 0.15 mM CaCl_2_, 10 mM K_2_HPO_4_/ KH_2_PO_4_ pH 7.6, 25 mM HEPES pH7.6, 2 mM EGTA, 5 mM MgCl_2_) using a BTX ECM 630 machine (Harvard Apparatus). Electroporation was performed in a 2 mm cuvette at 1.100V, 25 Ω and 25 µF. When needed, the antibiotic (concentration) used for drug selection was chloramphenicol (20 µM), mycophenolic acid (25 µg/ml) with xanthine (50 µg/ml), pyrimethamine (3 µM), or 5-fluorodeoxyuracil (10 µM). Stable transgenic parasites were selected with the appropriate antibiotic, single-cloned in 96 well plates by limiting dilution and verified by immunofluorescence assay or genomic analysis.

### Chromatographic purification of *Tg*PRP4K- and *Tg*PRP8-containing complex

*T. gondii* extracts from RHΔ*ku80* or PruΔ*ku80* cells stably expressing HAFlag-tagged *Tg*PRP4K and *Tg*PRP8, were incubated with anti-FLAG M2 affinity gel (Sigma-Aldrich) for 1 hour at 4°C. Beads were washed with 10-column volumes of BC500 buffer (20 mM Tris-HCl, pH 8.0, 500 mM KCl, 20% glycerol, 1 mM EDTA, 1 mM DTT, 0.5% NP-40, and protease inhibitors). Bound polypeptides were eluted stepwise with 250 μg/ml FLAG peptide (Sigma Aldrich) diluted in BC100 buffer. For size-exclusion chromatography, protein eluates were loaded onto a Superose 6 HR 10/30 column equilibrated with BC500. Flow rate was fixed at 0.35 ml/min, and 0.5-ml fractions were collected.

### Mass spectrometry-based Interactome analyses

Protein were stained with colloidal blue (Thermo Fisher Scientific) and gel bands excised before in-gel digestion using modified trypsin (Promega, sequencing grade). Resulting peptides were analyzed by online nanoliquid chromatography coupled to tandem MS (UltiMate 3000 RSLCnano and Q-Exactive HF, Thermo Scientific). Peptides were sampled on a 300 µm x 5 mm PepMap C18 precolumn and separated on a 75 µm x 250 mm C18 column (Reprosil-Pur 120 C18-AQ, 1.9 μm, Dr. Maisch) using 50-min gradients. MS and MS/MS data were acquired using Xcalibur (Thermo Scientific). Peptides and proteins were identified using Mascot (version 2.6) through concomitant searches against the *Toxoplasma gondii* database (ME49 taxonomy, version 30 downloaded from ToxoDB^47^, the Uniprot database (*Homo sapiens* taxonomy, February 2019 download), a homemade database containing the sequences of classical contaminants, and the corresponding reversed databases. Trypsin was chosen as the enzyme and two missed cleavages were allowed. Precursor and fragment mass error tolerances were set at respectively 10 ppm and 25 mmu. Peptide modifications allowed during the search were: Carbamidomethyl (C, fixed), Acetyl (Protein N-term, variable) and Oxidation (M, variable). The Proline software (http://proline.profiproteomics.fr) was used to filter the results: conservation of rank 1 peptide-spectrum-matches (PSMs), PSM homology threshold *p*-value ≤ 0.01, PSM score ≥ 25, and minimum of 1 specific peptide per identified protein group. Proline was then used to perform a compilation and grouping of the protein groups identified in the different samples. The MS data have been deposited to the ProteomeXchange Consortium via the PRIDE partner repository with the dataset identifier PXD029455 and 10.6019/PXD029455. Proteins from the contaminant database were discarded from the final list of identified proteins. MS1-based label free quantification of the protein groups was performed using Proline to infer intensity-based absolute quantification (iBAQ) values that were used to rank identified Toxoplasma proteins in the interactomes.

### Gene synthesis for recombinant expression of *Tg*PRP4K and *Pf*CLK3

Gene synthesis for all insect cell codon optimized constructs was provided by Genscript. The original *T. gondii Tg*PRP4K construct (aa 534-895) or *Pf*CLK3 (aa 336-692) were designed with non-cleavable C-terminal 6His tags and cloned between BamHI and HindIII sites into the pFastBac1 vector (Invitrogen). Point mutation variations of this initial construct were subsequently modified by Genscript from this original template. For the crystallization of *Tg*PRP4K, the cysteine 573 was mutated to a serine to prevent the formation of homomeric disulfide bond.

### Generation of baculovirus

Bacmid cloning steps and baculovirus generation were performed using EMBacY baculovirus (kindly gifted by Imre Berger), which contains a YFP reporter gene in the virus backbone. The established standard cloning and transfection protocols setup within the EMBL Grenoble eukaryotic expression facility were used. While baculovirus synthesis (V0) and amplification (to V1) were performed with SF21 cells cultured in SF900 III media (Life Technologies), large-scale expression cultures were performed with Hi-5 cells cultured in Express-Five media (Life Technologies) and infected with 0,5% vol/vol of generation 2 (V1) baculovirus suspensions and harvested 72h post-infection.

### Protein expression and purification

For purification, 3 cell pellets of approximately 800 mL of Hi-5 culture were each resuspended in 50 mL of lysis buffer (50 mM Tris pH 8.0, 500 mM NaCl and 4 mM β-mercaptoethanol (β-ME)) in the presence of an anti-protease cocktail (Complete EDTA free, Roche) and 1 ul benzonase (MERK Millipore 70746). Lysis was performed on ice by sonication for 3 min (30 sec on/ 30 sec off, 45° amplitude). After the lysis step, 5% of glycerol was added. Clarification was then performed by centrifugation for 1h at 12,000 xg and 4°C. After that, 20 mM imidazole was added to the supernatant and incubated with 5 mL of Ni-NTA resin (Qiagen) with a stirring magnet at 4°C for 30 min. All further purification steps were then performed at room temperature. After flowing through the lysate, the resin was washed with 10 column volumes of lysis buffer containing 20 mM imidazole. Elution was then performed by increasing the imidazole content to 300 mM in a buffer system containing 200 mM NaCl, 50 mM Tris pH 7.5, 2 mM BME and 5% glycerol. Eluted fractions were pooled based on an SDS PAGE gel analysis and flown directly through a previously equilibrated (in 200 mM NaCl, 50 mM Tris pH 7.5, 2 mM BME and 5% glycerol) heparin column connected to an AKTA© pure system. After a 10 cv wash, the heparin was eluted using a 40 mL gradient reaching 2M NaCl. Finally, the sample was injected onto a SUPERDEX 200 increase 10/300GL (GE Healthcare) column running in 50 mM Tris pH: 7.5, 250 mM NaCl, 1 mM BME and 1% glycerol and the elution was monitored at 280 nm. Peak fractions were concentrated using a 30 kDa Amicon ultra (Sigma Aldrich) concentrator before being frozen in liquid nitrogen and stored long-term at -80°C.

### Western blot

Immunoblot analysis of protein was performed as described in Farhat *et al.,* 2020. Briefly, ∼10^7^ cells were lysed in 50 μl lysis buffer (10 mM Tris-HCl, pH6.8, 0.5 % SDS [v/v], 10% glycerol [v/v], 1 mM EDTA and protease inhibitors cocktail) and sonicated. Proteins were separated by SDS-PAGE, transferred to a polyvinylidene fluoride membrane (Immobilon-P; EMD Millipore) by liquid transfer, and Western blots were probed using appropriate primary antibodies followed by alkaline phosphatase or horseradish peroxidase-conjugated goat secondary antibodies. Signals were detected using NBT-BCIP (Amresco) or enhanced chemiluminescence system (Thermo Scientific).

### Thermal Shift Assay (TSA)

The thermal stability of recombinant WT and mutants *Tg*PRP4K proteins in the presence or absence of altiratinib compound was performed in TSA buffer (400 mM NaCl, 50 mM Hepes, 1 mM MgCl_2_ and 2 mM beta-mercaptoethanol). Each assay tube contained a reaction mixture (final volume of 20µl) of recombinant PRP4K enzyme (0.170 mg/ml) and 100µM inhibitor or 1% dimethyl sulfoxide (DMSO) in TSA buffer. The reactions were incubated at increasing temperatures, 30°C, 33°C, 36 C°, 39°C, 42°C, 45°C, 48°C, 51°C, 54°C, 57°C, 60°C, 63°C, 66°C, 69°C, for 3 minutes and then centrifuged at 16000 xg for 25 minutes. The supernatant was collected to verify the presence of the recombinant PRP4K/CLK3 proteins by Western blot. Proteins were blotted as previously described and detected using Anti-polyHistidine-Peroxidase monoclonal antibody (Sigma-Aldrich # A7058) and signals were revealed using the Metal Enhanced DAB Substrate Kit, according to the manufacturer’s instructions (Thermo Scientific # 34065).

### Microscale thermophoresis (MST)

MST measurements were performed using a NanoTemper Monolith NT.115 Green/Red instrument (NanoTemper Technologies). *Tg*PRP4K protein was labeled using the Monolith His-Tag Labeling Kit RED-tris-NTA 2^nd^ Generation (NanoTemper Technologies). The labeled *Tg*PRP4K protein was adjusted to 100 nM with a buffer containing 30 mM Hepes (PH 7.5), 400 mM NaCl, 2% Glycerol, 0.5 mM beta-mercaptoethanol. A series of 16 1:1 dilutions of the ligands was prepared using the same buffer. Subsequently, each ligand dilution was mixed with 1 volume of the labeled *Tg*PRP4K protein. Samples were placed in Premium capillaries (NanoTemper technologies) for measurements. Instrument parameters were set to 40% LED power and 40% MST power. Data from three independently pipetted measurements were analyzed with MO. Affinity Analysis software (NanoTemper Technologies) using the signal from an MST-on time at 1.5 sec after T-jump.

### SEC-MALLS

All MALLS runs were performed using an S200 Increase SEC column (10/300 GL, GE Healthcare). Sample injection and buffer flow were controlled by a Hitachi L2130 pump. The SEC column was followed by an L-2400 UV detector (Hitachi), an Optilab T-rEX refractometer (Wyatt technologies), and a DAWN HELEOS-II multi-angle light scattering detector (Wyatt technologies). Injections of 50 μL were performed using protein samples concentrated at a minimum of 4 mg.mL−1, a constant flow rate of 0.5 mL.min−1 was used. Accurate MALLS mass prediction was performed with the Astra software (Wyatt Technologies). The curves were plotted using Graphpad (Prism).

### Crystallization with Altiratinib

For *Tg*PRP4K/Altiratinib co-crystal growth, *Tg*PRP4K at 2-5 mg/ml was incubated for 20 minutes with 400 μM of altiratinib prior to injection on an S200 column running 50 mM Tris pH 7.5, 250 mM NaCl, 1 mM B-ME and 1% glycerol. The eluted protein was then pooled and concentrated to 20 mg/ml. It should be noted that altiratinib generates a typical absorbance signature below 260 nm which increases with the concentration of the protein. Crystallization was setup using the hanging drop vapor diffusion method with *Tg*PRP4K/altiratinib mixed in a 1/1 ratio with 18% PEG 3350 and 0.18 M Potassium thiocyanate. Crystals appeared generally after 2 weeks. The crystals were harvested using Hampton nylon loops, cryo-protected in the mother liquor supplemented with 18-20% glycerol and flash frozen in liquid nitrogen.

### Data collection and structure determination

X-ray diffraction data for *Tg*PRP4k/altiratinib crystals were collected by the autonomous beamline MASSIF-1 at the European Synchrotron Radiation Facility (ESRF) beamline MASSIF-1 (Bowler et al., 2015; Svensson et al., 2015) using automatic protocols for the location and optimal centering of crystals (Svensson et al., 2018). Strategy calculations accounted for flux and crystal volume in the parameter prediction for complete datasets. Diffraction was performed at 100K. Data collection was performed using XDS (Kabsch, 2010) while amplitude scaling/merging was handled by the Staraniso server (Global phasing LTD). Molecular replacement solutions were obtained with Phaser (McCoy et al., 2007) (within Phenix) using the crystal structure of human PRPF4B bound to rebastinib [Protein Data Bank (PDB) code: 6CNH] as a template, the initial solution was then improved through cycles of manual adjustment in Coot (Emsley and Cowtan, 2004) and automated building in phenix autobuild (Terwilliger et al., 2008). The Altiratinib geometry restraints were generated in phenix using *elbow*. Refinement was performed using Refmac5, phenix resolve or Buster (Global Phasing Ltd). Final pdb model corrections were performed using the pdb-redo server.

### Structure representations

Structural representations of *Tg*PRP4K and *Pf*PRP4K/CLK3 were performed using UCSF-Chimera while the schematic representation of altiratinib interaction network was computed using Ligplot.

### Direct RNA sequencing by nanopore

The mRNA library preparation followed the SQK-RNA002 kit (Oxford Nanopore) recommended protocol, the only modification was the input mRNA quantity increased from 500 to 1000 ng, all other consumables and parameters were standard. Final yields were evaluated using the Qubit HS dsDNA kit (Thermofisher Q32851) with minimum RNA preps reaching at least 150 ng. For all conditions, sequencing was performed on FLO-MIN106 flow cells either using a MinION MK1C or MinION sequencer. All datasets were subsequently basecalled with a Guppy version higher than 5.0.1 with a Qscore cutoff > 7. Long read alignment were performed by Minimap2 as previously described (Farhat *et al.,* 2021). Alignments were converted and sorted using samtools.

### Differential splicing analysis

Splice correction, collapse, quantification and differential isoform representation was performed using the FLAIR pipeline (Tang et al., 2020) with standard parameters however keeping non-consistent isoforms after the correction stage. The difference splicing script was used to generate gtf track files and quantification histograms.

### Intron retention quantification

Prior to counting retained introns the original gff files from eupadDB were processed using the Agat-conv (https://github.com/NBISweden/AGAT) program “agat_convert_sp_gff2gtf.pl”. Introns were added to the gtf file with “agat_sp_add_introns.pl”. A per transcript intron retention ratio was calculated by counting per transcript intron counts divided by standard transcript counts (+1) using htseq-count with the following input parameters parameters:

- “htseq-count -f bam -r name -s yes -i Parent -t intron -m intersection-nonempty” for retained introns per-transcript
- “htseq-count -f bam -r name -s yes -i ID -t transcript -m intersection-nonempty” for total transcripts

Subsequent merging treatment of count data was carried out in excel worksheets. The intron retention analysis was limited to spliced genes with a minimum of 2 transcripts. Intron retention values above 1 were excluded as these values are probably the consequence of mis-annoted genes. Final distributions of retained intron ratios were done in GraphPad Prism.

### Software and Statistical analyses

Volcano plots, scatter plots, and histograms were generated with Prism 7. Sample sizes were not predetermined and chosen according to previous literature. All experiments were performed in biological replicates to allow for statistical analyses. No method of randomization was used. All experiments were performed in independent biological replicates as stated for each experiment in the manuscript. All corresponding treatment and control samples from RNA-seq were processed at the same time to minimize technical variation. Investigators were not blinded during the experiments. Experiments were performed in biological replicates and provided consistent statistically relevant results.

### *P. falciparum* EC50 determination

EC50 values were obtained as previously described (Nardella F *et al.,* 2020). Briefly, a range of 2-step serial dilutions of altiratinib (starting concentration 5 µM) and Dihydroartemisinin (DHA, starting concentration 50 nM) were used to assess the activity of the compounds. GraphPad Prism 8 was used to interpolate IC_50_ from three independent experiments run in triplicate. DHA and DMSO were used as positive and negative controls, respectively.

### *falciparum* treatment and harvesting for RNA extraction

3.10^9^ trophozoite-stage parasites aged of 24- to 30-hours post-red blood cell invasion were treated for 8 hours with a concentration of 2.5 µM of altiratinib (corresponding to the EC_90_) or 0.1% DMSO (vehicle). The concentration and incubation time chosen were checked previously for inducing no obvious growth phenotype. Parasites were then washed with RPMI (Gibco) and harvested using 0.15% saponin lysis of the surrounding red blood cell. After thorough washing with PBS, RNA was extracted using the RNeasy mini kit (Qiagen) and sent for analysis. RNA was harvested in two independent experiments.

## Data availability

Correspondence and requests for materials should be addressed to M.A.H. The Nanopore RNAseq data have been deposited in NCBI’s SRA data PRJNA774463. The coordinates and structure factors for the *Tg*PRP4k/altiratinib structure have been deposited in the PDB with the accession number 7Q4A. The mass spectrometry proteomics data were deposited with the ProteomeXchange Consortium via the PRIDE partner repository with the record identifier : pxd029455.

## Acknowledgments

This work was supported by the Laboratoire d’Excellence (LabEx) ParaFrap [ANR-11-LABX- 0024], the Agence Nationale pour la Recherche [Project HostQuest, ANR-18-CE15-0023, Project ApiNewDrug, ANR-21-CE35-0010-01, Project EpiKillMal, ANR-20-CE18-0006], the European Research Council [ERC Consolidator Grant N°614880 Hosting TOXO to M.A.H], and Fondation pour la Recherche Médicale [FRM Equipe # EQU202103012571] and Roux-Cantarini Fellowship attributed to FN. The proteomic experiments were partially supported by Agence Nationale de la Recherche under projects ProFI (Proteomics French Infrastructure, ANR-10-INBS-08) and GRAL, a program from the Chemistry Biology Health (CBH) Graduate School of University Grenoble Alpes (ANR-17-EURE-0003). The HTX Lab (EMBL Grenoble) are thanked for support in screening for crystal conditions and automatic mounting of crystals.

## Author Contributions

MAH led the research and coordinated the collaboration. MAH, AB and CS designed the project. CS, VB, MWB, FN, MPBP, CM, DC, YC, LB, FL, AB, AS and MAH designed, conducted and interpreted the experimental work. M.-A.H., A.B. and C.S. wrote the paper with editorial assistance from VB and fruitful comments from all other authors.

---

**Extended Data Fig. 1.**
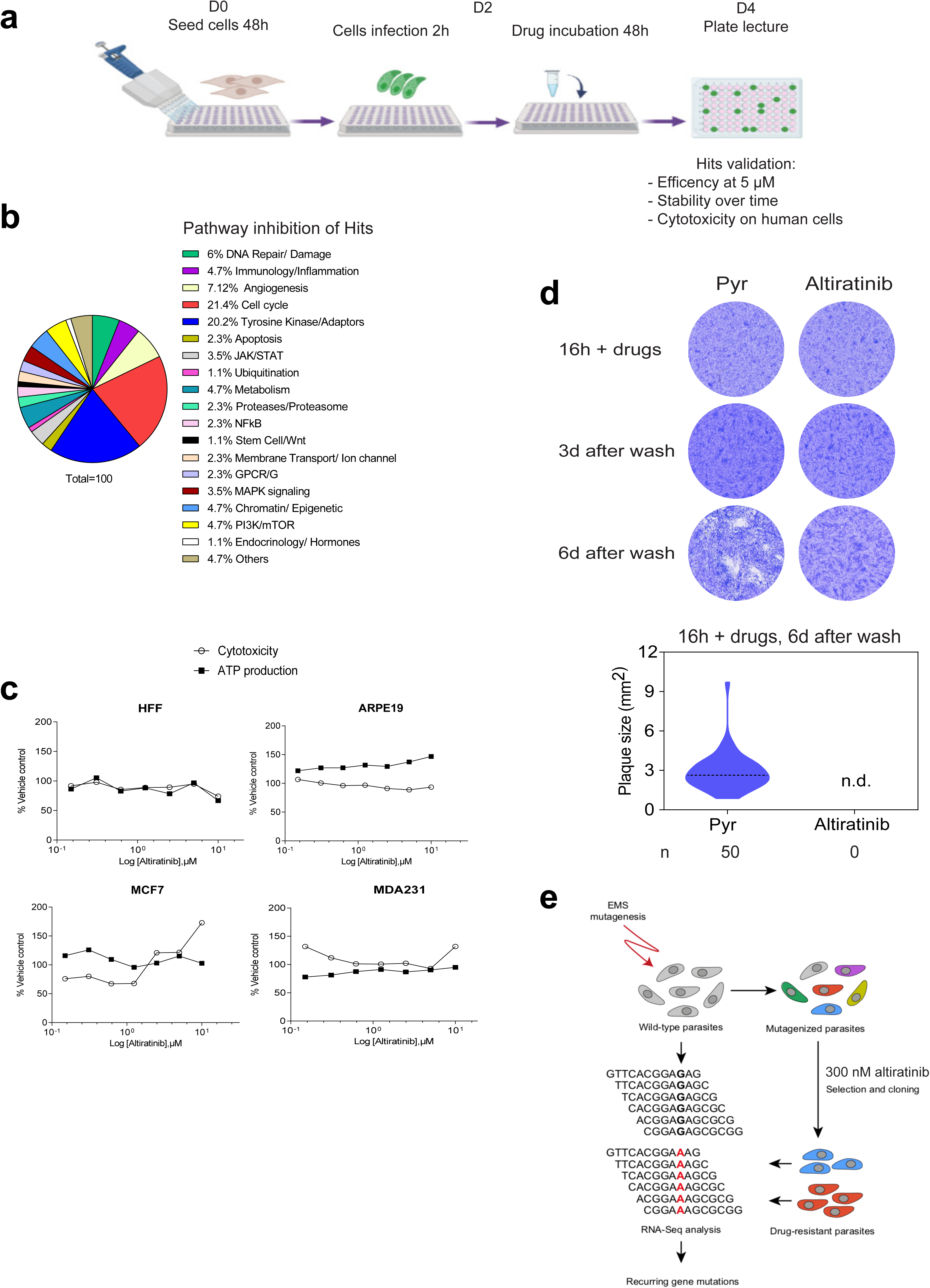
Identification of altiratinib by a medium-throughput screening of an FDA-approved library. **a,** Schematic overview of the workflow used to screen the library of514-FDA approved compounds. Confluent HFFs were infected for 2 hours with tachyzoites of the *T. gondii* RH strain expressing the NanoLuc luciferase (RH Δ*ku80 UPRT::NLuc-P2A-EmGFP*). Each compound was then added to the culture at a concentration of 5 µM for 48h. After washing and incubation with furimazine substrate, luciferase activity was detected to select hits. Hits were further validated by testing their efficiency at 1 µM and checking stability over time or toxicity to the host cells. **b,** Distribution of 84 hits by pathway inhibition. **c,** Mitochondrial toxicity assay. Human cells were incubated with increasing concentrations of altiratinib, sodium azide (positive control for mitochondrial toxicity) or 800µg/ml of digitonin (positive control for cell toxicity). After 90 min, cell viability (cytotoxicity) was detected by fluorescence readout, while ATP production was measured by luminescence as indicated in the “*Mitochondrial ToxGlo^TM^ Assay”* kit (Promega). Plots are representative of two biological replicates. **d,** Representation of *T. gondii* cytotoxicity after incubation with drugs. Confluent HFFs were infected with the strain RH WT (RH Δ*ku80 UPRT::NLuc-P2A-EmGFP*) and incubated with 1 µM of pyrimethamine or 300nM of altiratinib for 16h. After 3 and 6 days, the drugs were washed out and the cells were stained with Coomassie blue to detect the presence of plaques. Graphs show the size of visible plaques in each condition. n.d., not detected. Statistical analyses were performed using Mann-Whitney test (One-way ANOVA). **e,** Workflow used to map mutations that confer resistance to altiratinib in parasites.

**Extended Data Fig. 2.**
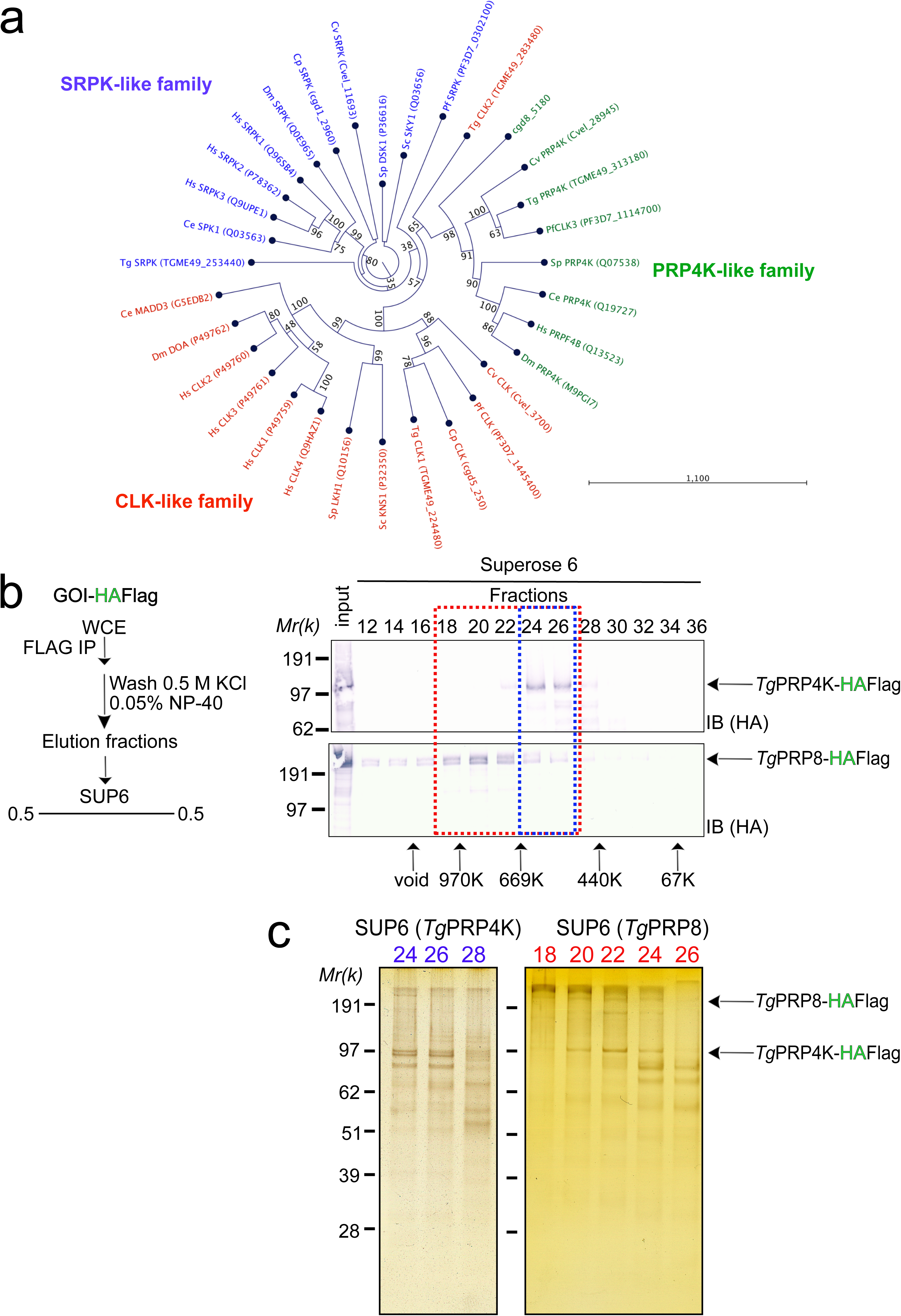
Origin and interactome of *Tg*PRP4K. **a,** Phylogenetic analysis of CLK-, PRP4K- and SRKP-like families. The unrooted phylogenetic tree was inferred from the kinase domain alignment. The tree was computed with the neighbor-joining algorithm, based on an HMM multiple alignment. The bootstrap values are shown in blue. The reliability of branching was assessed by the bootstrap resampling method using 1000 bootstrap replicates. **b and c,** Size-exclusion chromatography of PRP4K- and PRP8-containing complexes after Flag-affinity selection. The fractions were analysed using western blots to detect PRP4K- or PRP8– HAFlag (anti-HA antibodies) (**b**) and on silver-stained SDS–PAGE gels (**c**).

**Extended Data Fig. 3.**
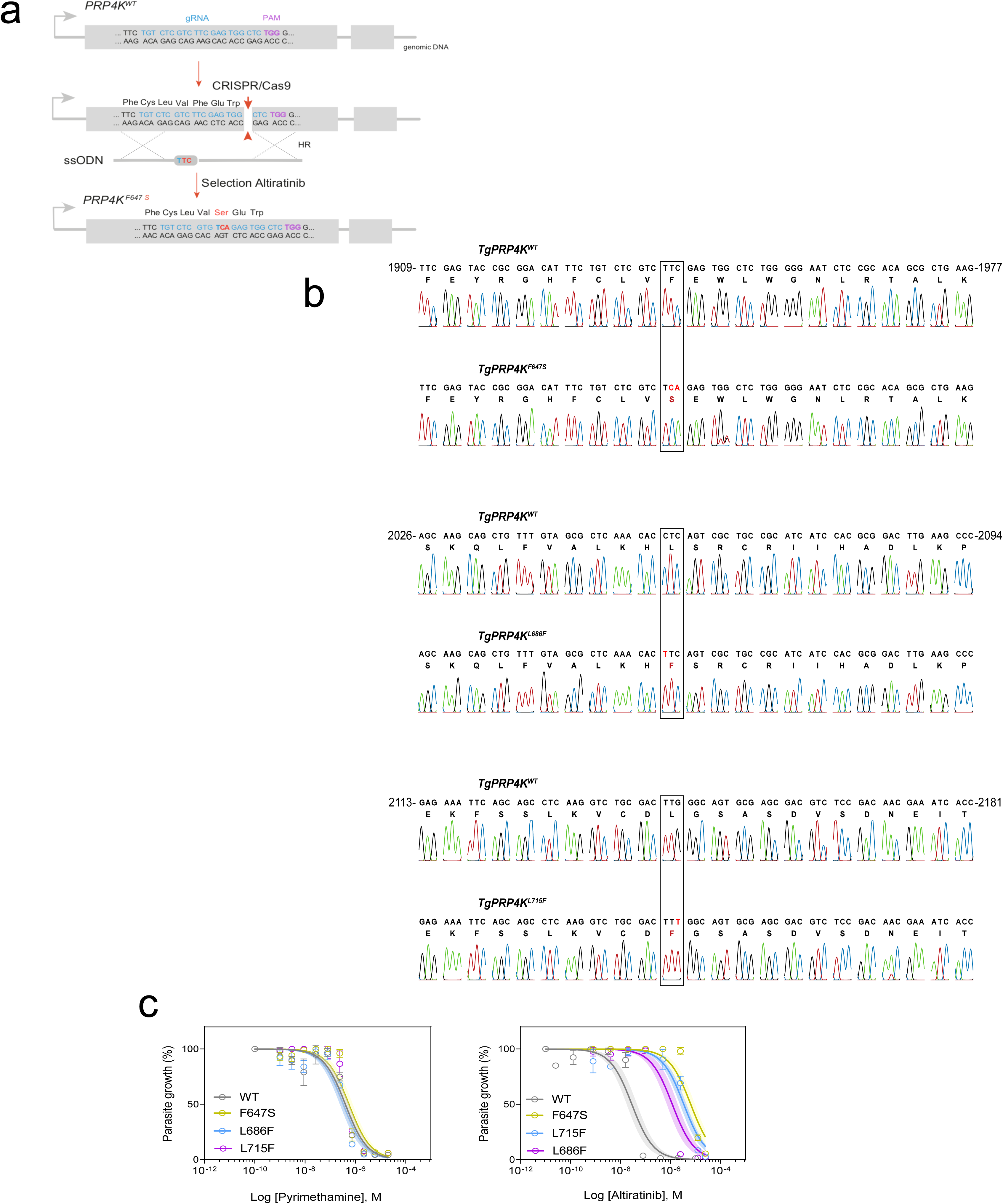
Identification and validation of the molecular target *Tg*PRP4K. **a,** Strategy for introducing point mutations into *T. gondii* parasites. Focus on the *Tg*PRP4K locus and CRISPR/Cas9-mediated homology-directed repair with single-stranded oligo DNA nucleotides (ssODNs) carrying nucleotide substitutions (red letters). After homologous recombination (HR) events, the *Tg*PRP4K recombinant parasites were selected with altiratinib. The engineered parasites were then validated by Sanger sequencing. **b,** Sanger chromatograms validating *TgPRP4K* gene editing. Indicated are the nucleotide positions relative to the ATG start codon on the genomic DNA. **c,** Dose–response curves for inhibition of *T. gondii* growth in response to increasing concentration of the indicated compounds. Confluent HFF monolayer were infected with WT and the engineered *Tg*PRP4K mutant strains expressing the NanoLuc luciferase. Data are presented as mean ± standard deviation (SD) of n=3 technical replicates from a representative experiment out of at least two independent biological assays. Shaded error envelopes depict 95% confidence intervals.

**Extended Data Fig. 4.**
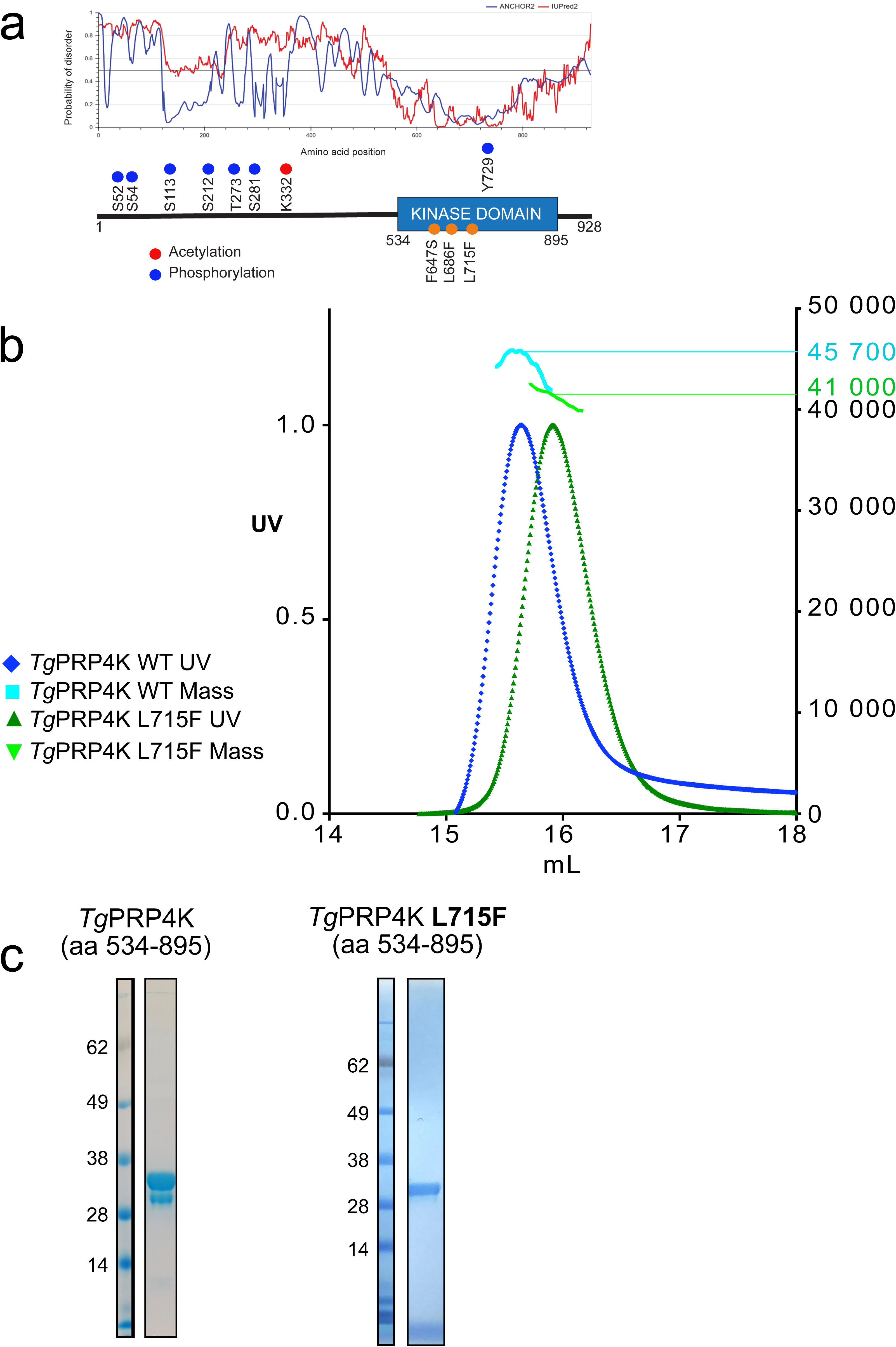
Insect-cell recombinant expression of *Tg*PRP4K. **a,** PRP4K full protein organization. IUpred disorder propensity prediction is shown above a linear schematic representation of the polypeptide chain. Phosphorylated and Acetylated residues (data extracted from ToxoDB.org) are highlighted in blue and red respectively. **b,** SEC-MALLS measurement of *Tg*PRP4K sample homogeneity. UV(280nm) absorbance chromatogram of insect cell purified WT (in blue) or L715F mutant (in green) *Tg*PRP4K (534-895) combined to a mass calculation as a scatter plot with the Y axis (in KDa) on the right when run on a S200 increase column. **c,** SDS PAGE gel of recombinantly produced PRP4K. WT (left) and L715F (right) purified protein analysed a NuPage 5-12% Bis-Tris gel run in MES buffer and colored with Coomassie blue. The indicated numbers correspond to protein marker mass in kDa.

**Extended Data Fig. 5.**
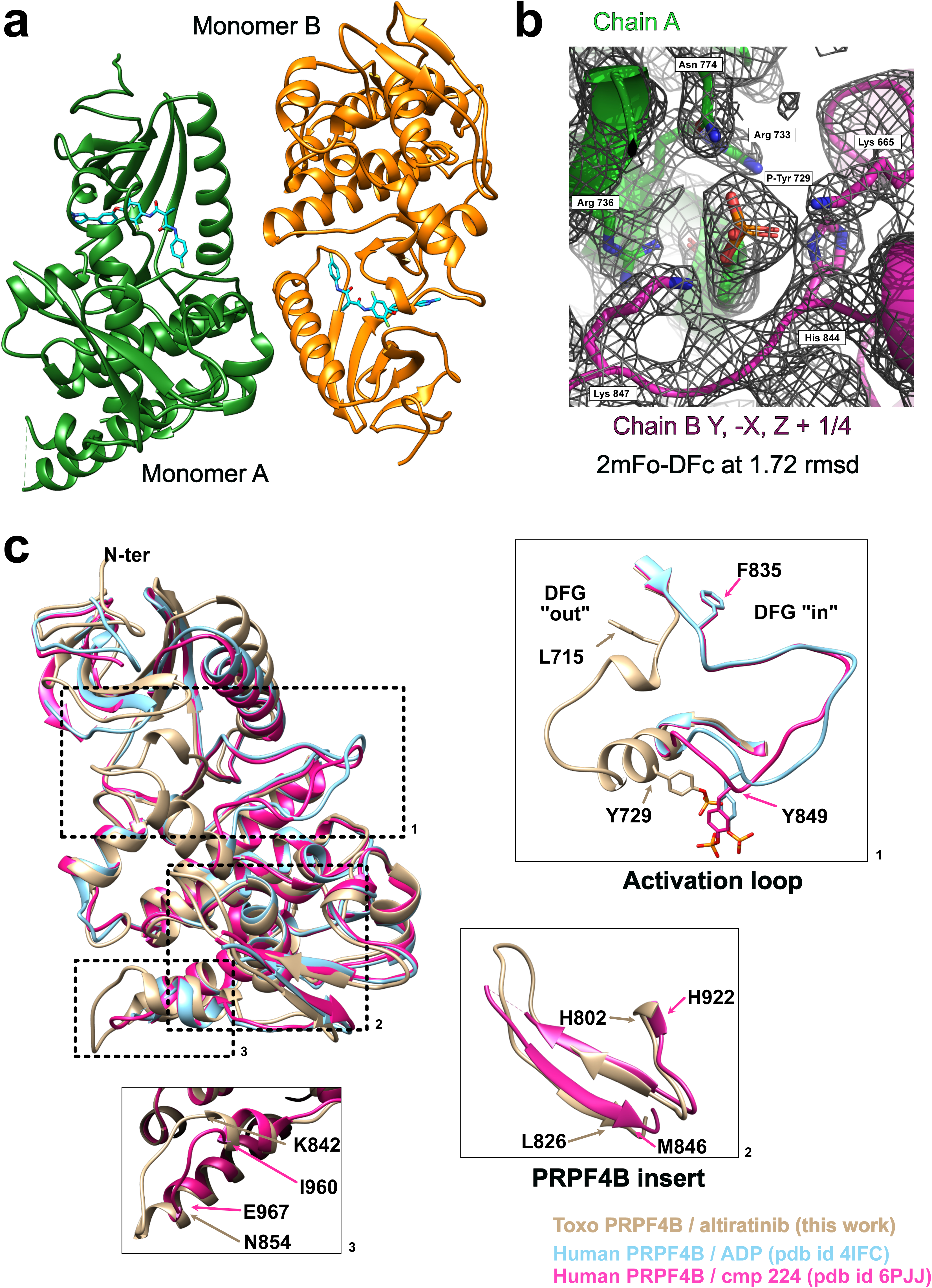
Crystal structure specificities of *Tg*PRP4K. **a,** Crystallographic dimerization of *Tg*PRP4K. Cartoon representation of chain/monomer A and B colored green and orange respectively. Altiratinib is displayed as sticks in cyan. **b,** Phosphotyrosine 729 mediated crystal contacts. Cartoon and stick representation of the phosphotyrosine and interaction side chains from homo-monomeric and symmetry related molecules. 2mFo-DFc electron density is represented at an rmsd of 1.72. This specific representation was done using pymol. **c,** Structural conservation of *Tg*PRP4K compared to the human *Hs*PRPF4B. Structural alignment shown in a cartoon fashion between *Tg*PRP4K bound to altiratinib (tan), ADP bound *Hs*PRPF4B (skye blue) and *Hs*PRPF4B (magenta) bound to cmp 224. Certain regions are shown enlarged for more detail.

**Extended Data Fig. 6.**
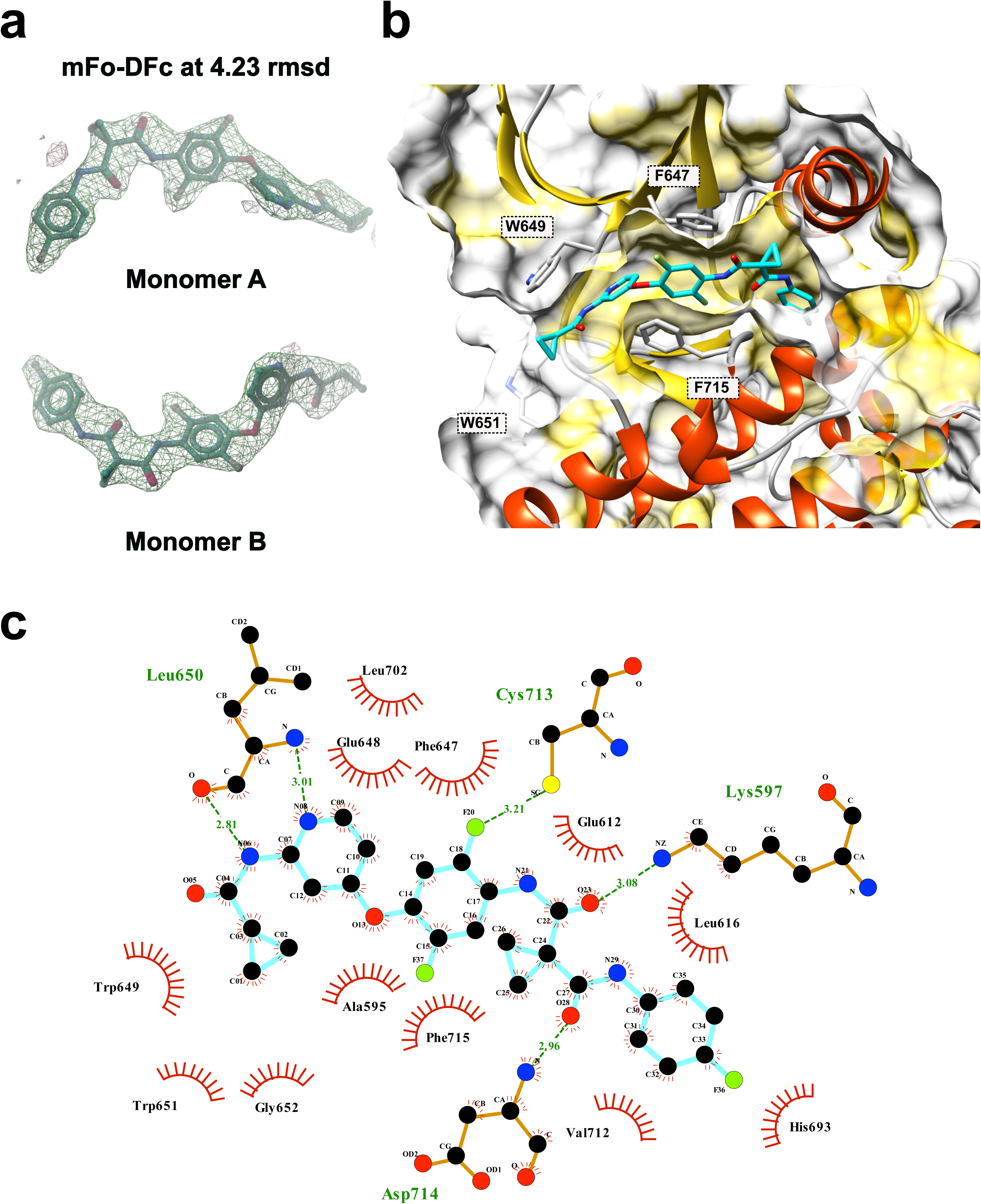
Altiratinib binding site and interaction network. **a,** Altiratinib omit map. Altiratinib mFo-DFc omit map (generated in coot) at 4.2 rmsd showing electron density as a green/grey mesh and the altiratinib stick structure in green. **b,** ATP and allosteric pocket binding of altiratinib. Cartoon representation combined with a surface and hydrophobic attribute coloring (in yellow). Notable side chains involved in hydrophobic interactions are displayed as stick side chains. **c,** Ligplot schematic 2D representation of all residues interacting with altiratinib. Charged interactions are displayed as green dotted lines whereas hydrophobic interactions are shown with red curved-in combs.

**Extended Data Fig. 7.**
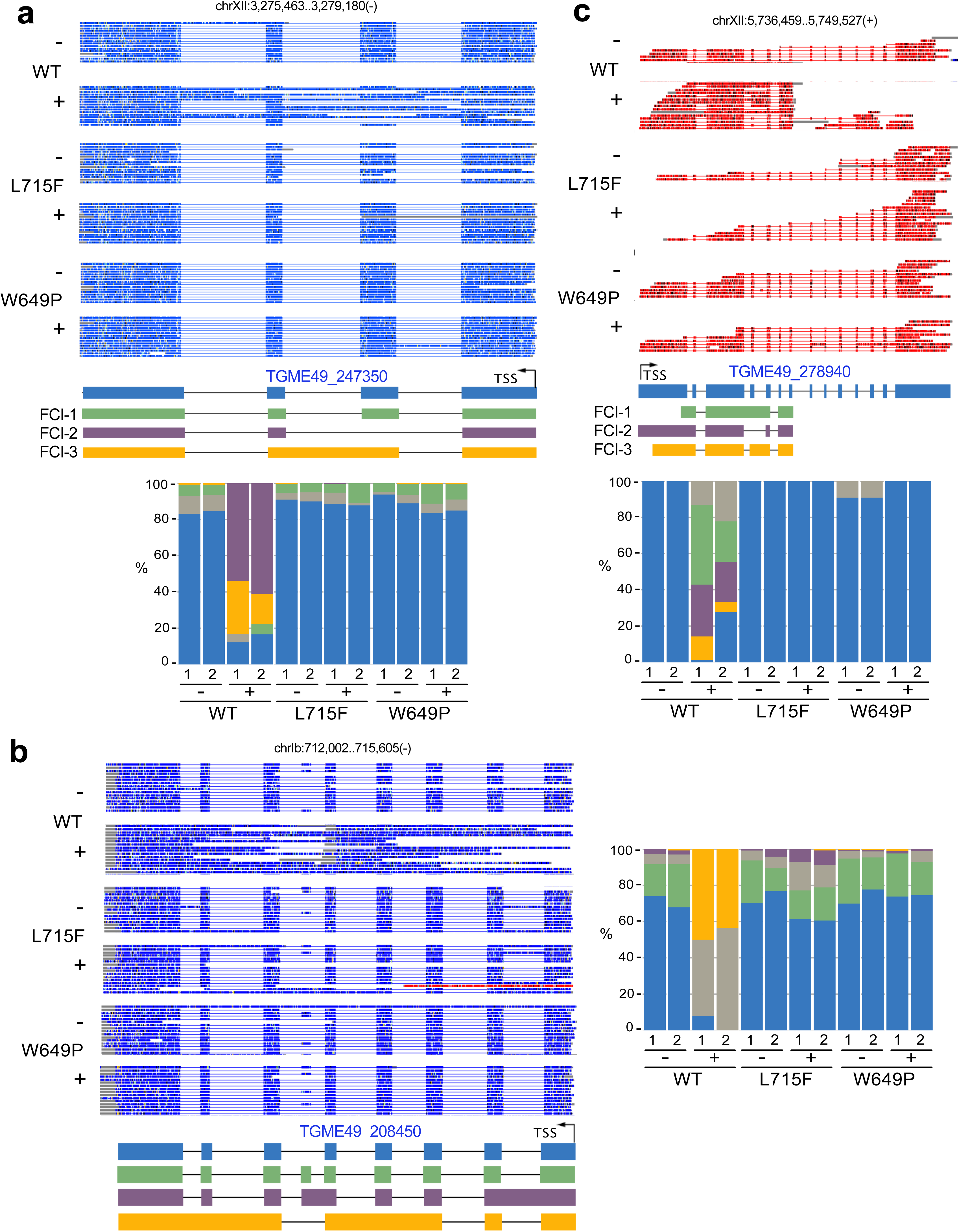
DRS examples of altiratinib induced splicing defects in *Toxoplasma gondii*. FLAIR analysis of *TGME49_247350* (**a**), TGME49_208450 (**b**) and *TGME49_278940* (**c**) loci. Standard annotation and FLAIR collapsed isoforms (FCI) are shown schematically for all panels under a sample view of 15 reads per condition. Sense and antisense reads are colored red and blue, respectively. Below the FCI representation is an isoform quantification histogram showing duplicate measurements in each WT/L715F/W649P and untreated (-) or treated (+) condition. The color code is the same as for the above FCI, grey histograms represent minor isoforms that are not schematically represented.

**Extended Data Fig. 8.**
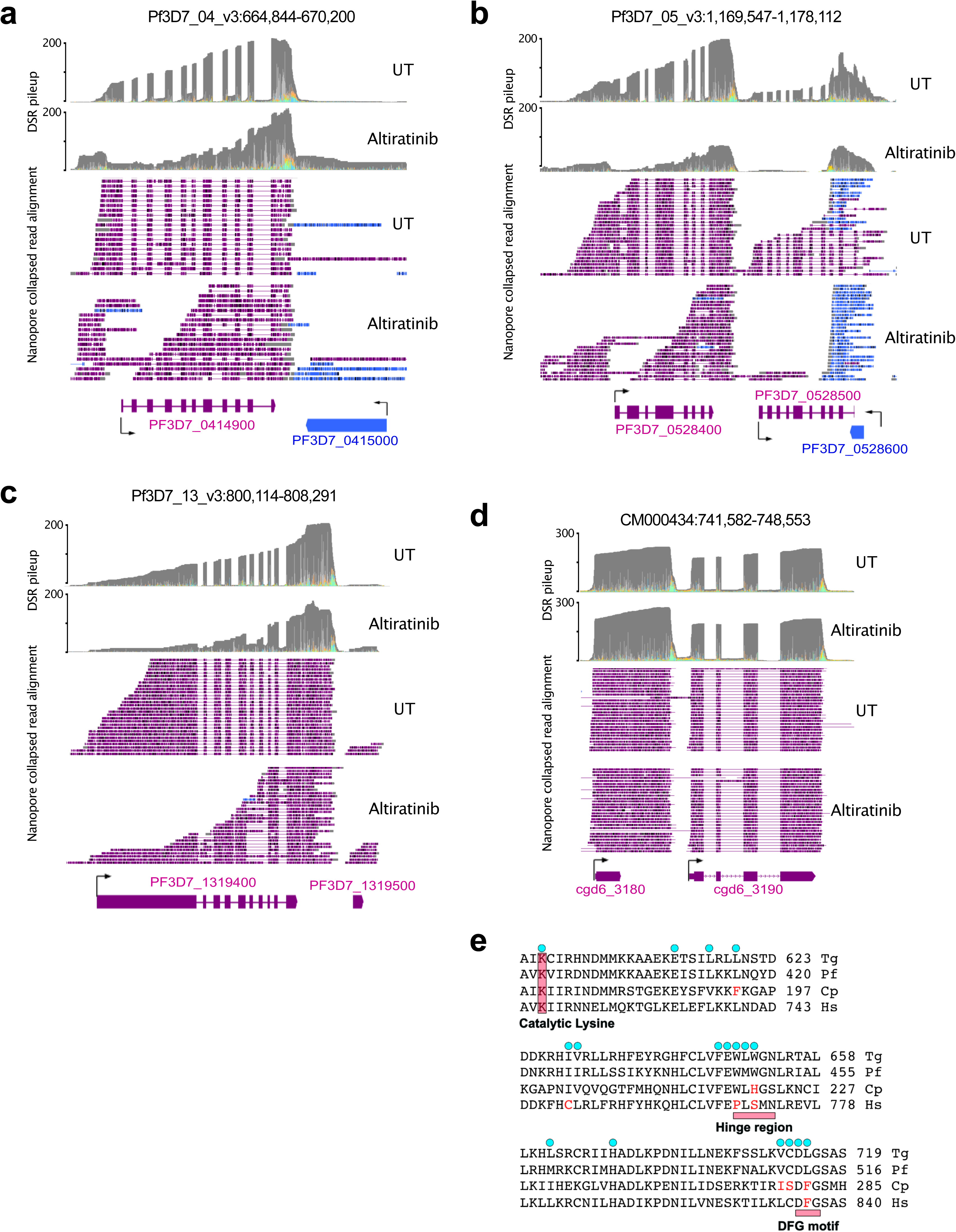
DRS examples of altiratinib treatment on *Plasmodium falciparum* and *Cryptosporidium parvum*. FLAIR analysis of *PF3D7_0414900* (**a**), *PF3D7_0528400* (**b**), *PF3D7_1319400* (**c**) and *PF3D7_* loci in *Plasmodium falciparum* as well as the *cgd6_3190* loci (**d**) in *Cryptosporidium parvum*. Standard annotation and FLAIR collapsed isoforms (FCI) are shown schematically for all panels under a sample view of 15 reads per condition. Sense and antisense reads are colored purple and blue, respectively. Below the FCI representation is an isoform quantification histogram showing duplicate measurements in untreated (-) or treated (+) condition. The color code is the same as for the above FCI, grey histograms represent minor isoforms that are not schematically represented. **e,** Sequence alignment of the *Tg*PRP4K binding regions of altiratinib compared to PRP4K/CLK3 orthologs of *Plasmodium falciparum* (*Pf*), *Cryptosporidium parvum* (*Cp*) and *Homo sapiens* (*Hs*). Key regions are highlighted by pink rectangles, altiratinib interacting amino acids from *Tg*PRP4K are shown by cyan circles, while divergent residues at the same position in the orthologs *of Cp* and *Hs* are shown in red.

**Extended Data Fig. 9.**
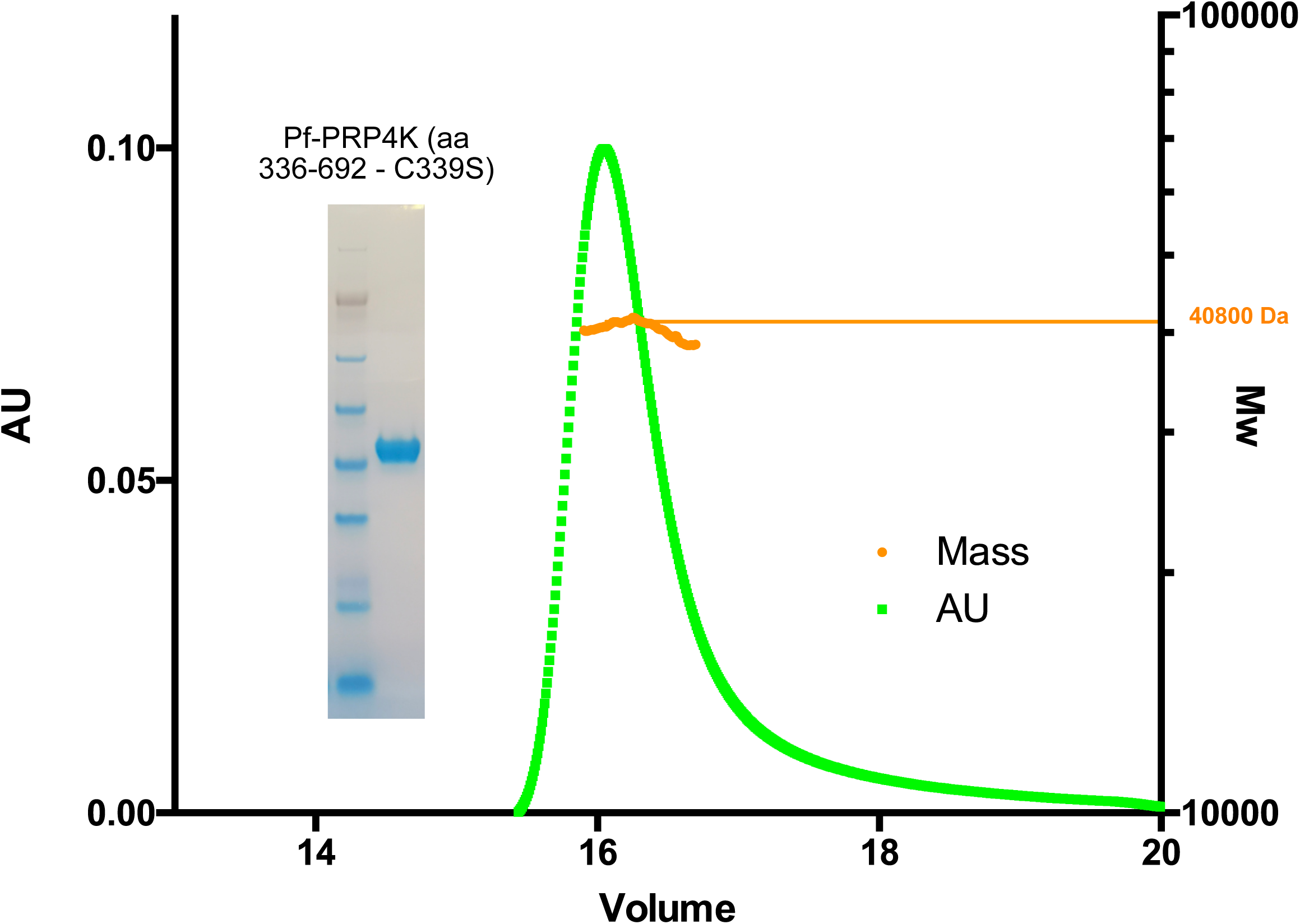
Biochemistry of recombinant *Pf*PRP4K. SEC-MALLS chromatogram of homogeneity of *Pf*PRP4K sample. UV (280 nm) absorbance chromatogram of insect cell-purified *Pf*PRP4K (aa 336-692 with the C339 mutated to a S) combined to a mass calculation as a scatter plot with the Y axis (in KDa) on the right when run on a S200 increase column. Next to the chromatogram is a NuPAGE 4-12% gel of the same purified sample run in MES and stained with Coomassie blue. The numbers shown correspond to the marker mass in kDa.

## Supplementary files

**Supplementary Fig. 1.**
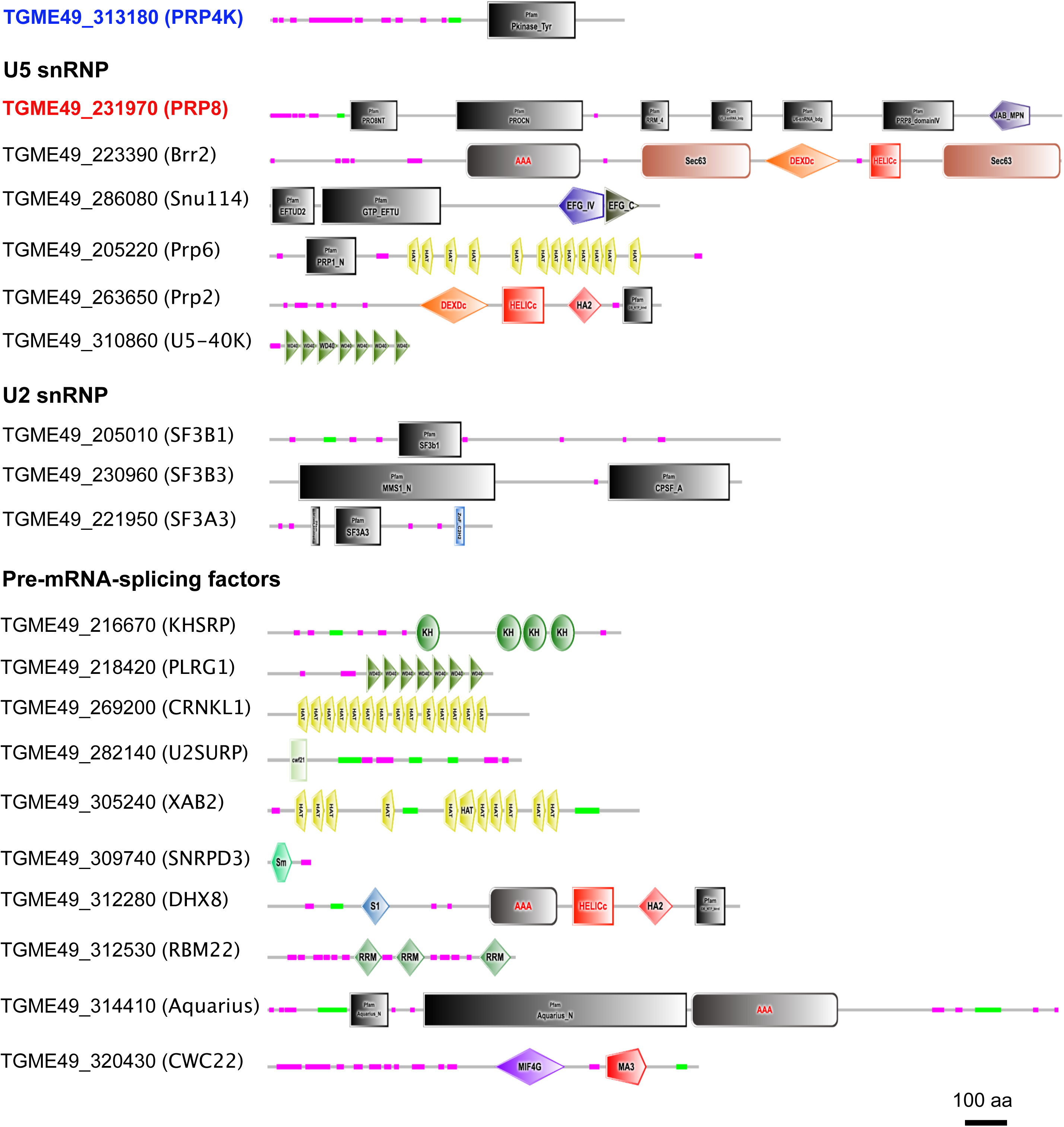
Domain architectures of proteins purified together with PRP4K and PRP8. Representative domain architectures of T. gondii PRP4K and PRP8 partners identified by mass spectrometry-based proteomics (Supplementary Table 3) are shown. Domains were predicted by SMART and PFAM.

**Supplementary Table 1 | Table describing the 514 compounds in the TargetMol© FDA-approved compound library.**

**Supplementary Table 2 | Mutations found in candidate genes by RNA-sequencing analysis of altiratinib-resistant mutants.** Amino acid substitutions with corresponding codons shown in parentheses are indicated for each *T. gondii*-resistant mutant strain.

**Supplementary Table 3 | Mass spectrometry-based characterization of the interactomes of PRP4K and PRP8.** *Tg*PRP4K- and *Tg*PRP8-containing complexes were purified by affinity using FLAG tagging and subjected to size exclusion chromatography. Proteins present in the different eluted fractions were pooled and separated by SDS-PAGE. For *Tg*PRP4K, two lanes were analyzed with pools of fractions 24 and 26, and fractions 28 and 30. For *Tg*PRP8, three lanes were analyzed with fractions 14, 16 and 18, fractions 20 and 22, and fractions 24 and 26 (see below). The bands of interest in each lane, marked with arrows below, were excised and analyzed using MS-based proteomics. The parasite proteins identified and quantified in each band of the *Tg*PRP4K- and *Tg*PRP8-containing complexes are listed in the following tables.

**Supplementary Table 4 | Statistics of crystallographic data.**

**Supplementary Table 5 | Description of T. gondii strains, plasmids and primers.** List of *T. gondii* parasite lineages as well as plasmids used in this work. Primers used in this work are also listed.

**Supplementary Data File 1 | Full PDB X-ray structure validation report** of crystal structure of *Toxoplasma* TgPRP4K with altiratinib (pdb id: 7Q4A).

